# Adaptive Strategies of the invasive aquatic plant, *Ludwigia grandiflora* subps. *hexapetala*: Contrasting Plasticity Between Aquatic and Terrestrial Morphotypes

**DOI:** 10.1101/2025.08.27.672630

**Authors:** Julien Genitoni, Danièle Vassaux, David Renault, Stéphane Maury, Dominique Barloy

## Abstract

Biological invasion is the fifth biggest threat to biodiversity and ecosystem processes. Invasive species are able to colonise new environments rapidly due to their phenotypic plasticity. *Ludwigia grandiflora* subsp. *hexapetala* (*Lgh*), an aquatic invasive plant, invades various aquatic habitats, such as rivers, ponds, and more recently, especially in France, wet meadows. More recently, a terrestrial morphotype, emergent for half the year, has been identified. The behaviour of aquatic and terrestrial morphotypes in aquatic and terrestrial conditions was analysed through observations of morphological traits, and assays of metabolomics and phytohormones 14 and 28 days after the beginning of the experiment. The phenotypic plasticity was evaluated by calculating the relative distance plasticity index (RDPI) in response to environmental changes (terrestrial *versus* aquatic). RDPI were measured for morphological traits, metabolic and phytohormonal contents. Our results revealed that the terrestrial morphotype showed higher morphological trait values than the aquatic morphotype, independently of conditions, suggesting a pre-adaptation of *Lgh* to terrestrial habitats. The aquatic and terrestrial conditions shaped the *Lgh* metabolomic responses. However, the aquatic morphotype displayed higher phenotypic plasticity indexes than the terrestrial one. In addition, plasticity indexes evolved during the acclimatisation process, leading to increased RPDI values in most cases. The two morphotypes present distinct responses to aquatic and terrestrial conditions (especially in aquatic conditions for the terrestrial morphotype), highlighting the capacity of *Lgh* for invading new habitats due to its phenotypic plasticity, and a potential local adaptation. This study contributed to the understanding of how invasive species mobilise their phenotypic plasticity in new habitats, and the time required for local adaptation to occur.

## Introduction

During the invasion process, several determinants, including phenotypic plasticity, contribute to the success of the introduced propagules (Li, Du, Guan, Yu, & van Kleunen, 2016; Oduor, Leimu, & van Kleunen, 2016; Szabó, Peeters, Várbíró, Borics, & Lukács, 2019, Vedder et al, 2021). Lande (2015) highlights that adaptation to a new environment can be done through a transient increase in plasticity, followed by a decrease in plasticity involving slow genetic assimilation. Thus, it seems to be pertinent to monitor changes in plasticity after the introduction in order to assess the role of plasticity in the invasion success (Lande, 2015; Murren *et al*., 2015).

Native to South America, *Ludwigia grandiflora* subsp*. hexapetala* (named *Lgh)* is now invasive in the United States, Japan, Australia, Switzerland, the UK, and in many mainland European countries (France, Ireland, Spain, the Netherlands, Germany, Italy, Belgium) (Delange et al., 2025; Hussner et al., 2016; Pelella & and Ceschin, 2024). In Europe, *Lgh* is considered to be one of the most invasive aquatic plants (Hussner, Windhaus, & Starfinger, 2016). This species arrived in France in the 19^th^ century, and has rapidly colonised static or slow-flowing waters, river banks, and more recently wet meadows (Dandelot et al., 2005; Lambert et al., 2010). *Lgh* is a heterophyllous amphibious plant, able to produce pneumatophores and tolerant to submergence (Kuwabara et al., 2003). These characteristics suggest that *Lgh* displays phenotypic plasticity in changing environments (Wells & Pigliucci, 2000; Kuwabara & Nagata, 2006; Nakayama, Sinha, & Kimura, 2017).

For floating plant species, the transition from aquatic to terrestrial environments requires the development of numerous adaptation strategies in order to cope with the temporal terrestrial habitat (Li Z, Yu D, Xu J, 2011; Robe & Griffiths, 1998). *Lgh* shows two distinct morphotypes in function of the invaded habitat (aquatic or terrestrial) (Billet et al., 2018; Haury et al., 2014). One of the two morphotypes is considered as aquatic (aquatic morphotype) and spends its entire life cycle in water. The second is considered to be terrestrial (terrestrial morphotype) and grows in wet meadows, with seasonal water level fluctuations (underwater from November to April and emergent the rest of the year). Haury et al., (2014) and Billet et al., (2018) worked in the field and in the growth chamber, respectively, and demonstrated the morphological differences between aquatic and terrestrial morphotypes. In particular, the aquatic morphotype had a lower number of nodes, a smaller number of leaves, and a higher water content than the terrestrial morphotype. The terrestrial morphotype, on the other hand, presented a greater number of leaves and a higher root mass (Haury et al., 2014; Billet et al., 2018).

Aquatic and terrestrial environments impose specific constraints on *Lgh* morphotypes. The aquatic condition is a carrier medium, favouring plant flotation and altering the access to oxygen and light, potentially leading to hypoxic stress. The shift from an underwater environment (aquatic environment) to an air environment with higher light and oxygen levels (terrestrial environment) can lead to drought stress (Tamang & Fukao, 2015). Thus, fluctuating water levels, which commonly occur in wetlands and rivers, can influence growth, plant length and resource allocations (Areington et al., 2024; Deegan et al., 2007; Rea & Ganf, 1994). When submerged, both the decrease in oxygen availability and loss of oxygen production by photosynthesis leads plants to shift metabolism from aerobic respiration to fermentation (Voesenek & Bailey-Serres, 2015). Metabolism pathways implicating sugars, amino acids or polyamines and phytohormones are often involved in drought and flooding stress responses (Hu et al., 2016; Loreti et al., 2016, 2018). Billet et al., (2018) identified specific metabolite patterns in *Lgh* for shoots growing in aquatic conditions, roots growing in terrestrial conditions, and the respective amino acid and sugar metabolomic pathways.

In this study, we worked on a sympatric population of two morphotypes from the marshlands of Mazerolles (near Nantes, France). The aquatic morphotype appeared here in 1995 and the terrestrial morphotype developed five years later. The five-year period between the arrival of the aquatic form and the transition to the terrestrial environment raises the question of the rapid adaptation of *Lgh* to a new habitat and the fate of the invasion (Billet et al., 2018). To investigate whether the fast transition to the terrestrial habitat changes *Lgh*’s physiological responses, and to explore how the phenotypic plasticity is affected, we compared the behaviour of aquatic and terrestrial morphotypes in both their original conditions and in changing environment (aquatic *versus* terrestrial), in controlled conditions. We characterised the morphological, phytohormone, and metabolomic responses in function of the condition 14 and 28 days after the beginning of experiment. We evaluated the phenotypic plasticity using the Relative distance plasticity index (RDPI) recommended by Valladares, F., Sanchez-Gomez, D., & Zavala, M. A., (2006), and followed its evolution over the duration of the experiment. We tested the following three hypotheses: (1) *Lgh* can adjust its morphology and physiology depending on the allocation of resources and will mobilise specific metabolomic pathways according to the habitat (Billet et al., 2018); (2) As the terrestrial morphotype undergoes consecutive submergence and desubmergence conditions, its phenotypic plasticity could be greater than those of the aquatic morphotype (Murren et al., 2015); (3) if *Lgh* expresses an adaptative phenotypic plasticity, the plasticity of the terrestrial and aquatic morphotypes in the reverse condition will evolve over time, eventually resembling that of the morphotype in its original condition (Ghalambor et al., 2007).

## Materials and methods

### Plant material and experimental design

Aquatic (Am) and terrestrial (Tm) morphotypes of *L. grandiflora* subsp. *hexapetala* were collected from the marshlands of Mazerolles (near Nantes, France, N47 23.260 W1 28.206). Following collection, plants were grown in a growth chamber in their initial conditions, i.e., aquatic conditions with a high-water level for Am, and terrestrial conditions with a low level of water for Tm at 22 °C, 16h/8h light/darkness cycle as described in Billet et al., (2018).

Stem cuttings of 10cm were collected from the stem apex, without roots, buds or lateral stems. Next, stems underwent a two-week preconditioning period in a growth chamber to promote the development of roots, as described in Billet et al., (2018). Then, for both aquatic and terrestrial morphotypes, plants for morphological and metabolomics analyses were randomly placed into containers (Length x Width x Height: 8cm x 8cm x 15cm) in either aquatic and terrestrial conditions, under the same controlled conditions. Am and Tm were placed in their original conditions (aquatic for Am (named Am-a) and terrestrial for Tm (named Tm-t), and in their reverse conditions (terrestrial for Am (named Am-t) and aquatic for Tm (named Tm-a) (Fig. S1).

Measurements of morphological variables and metabolic/phytohormone dosages were conducted two (t14) and four (t28) weeks after the transfer into containers. Two and three biological replicates were carried out for the morphological and metabolomic/phytohormones analyses, respectively. For metabolomic and phytohormone analyses, samples corresponding to the first ten cm of shoot apex were pooled and immediately snap-frozen in liquid nitrogen, then lyophilized over 48 hr using a Cosmos 20K (Cryotec, Saint-Gély-du-Fesc, France) and stored at −80 °C.

### Morphological analyses

Twelve plants per morphotype, condition, and biological replicate were analysed in order to investigate the effects of our experimental treatments on the growth and fitness traits of *Lgh*. Shoot length of the plant (LP, in centimeter), leaf number (nbL) and inter-node number (nbI), which characterised plant morphology, were noted. These data were used to analyse the growth strategy of *Lgh* in different conditions by calculating the nbL/Lp (NbLLP) and nbI/Lp (NbILP) ratios. *Lgh* can form new plants from small (>1 cm) plant fragments (Hussner, 2009).The ability to produce propagules from buds or nodes with roots can be considered a plant fitness trait, and can thereby be assessed by counting the number of nodes with buds (nbNB). Biomass of plants in grams (g) was determined by measuring the fresh mass of shoots (FMS) and roots (FMR). After oven drying the samples at 105 °C for 48 hr, we obtained their respective dry masses (DMS and DMR) and total dry mass (TDM=DMS+DMR) was calculated. To determine the distribution of water content in roots and shoots of *Lgh* morphotypes, the ratios of fresh/dry mass of shoots (Sr =FMS/DMS) or roots (Rr=FMR/DMR) were calculated. To understand the resource allocation of Am and Tm morphotypes according to the conditions analysed, ratios of dry mass roots/total dry mass (AlocR) and dry mass shoots/total dry mass (AlocS) were calculated.

### Metabolomic fingerprint

Six plants per morphotype, per experimental condition and per biological replicate were sampled at 14 and 28 days, as described in the section “Plant material and experimental design”. We used a gas-chromatograph coupled to a mass-spectrometer (GC-MS), and the methodology described in Serra et al. (2013) and adapted by (Billet et al., 2018b). For each sample, an aliquot of 10 mg of lyophilized powder was homogenized into 600 μL of a solution of ice-cold methanol/chloroform (2:1, v/v). Then, a volume of 400 μL of ultra-pure water was added. Samples were homogenized and centrifuged for 10 min at 4,000 g (4 °C). Next, 120 µL or 80 µL of the upper phase containing metabolites were transferred to new glass vials for roots and shoots, respectively. Chromatograms were analysed with XCalibur 2.0.7. The concentration of each metabolite was calculated using individual quadratic calibration curves.

### Phytohormone quantification

Quantification of abscisic acid (ABA), salicylic acid (SA) and auxin (IAA) were carried out for shoots at 28 days. Each sample was constituted by six plants per morphotype, per experimental condition and per biological replicate. For each sample, 1 mg of dry powder was extracted with 0.8 mL of acetone/water/acetic acid (80/19/1 v/v/v). Each dry extract was then treated as described in Genitoni et al., (2020).

### Phenotypic plasticity

The phenotypic plasticity of the plants exposed to the different experimental conditions was calculated by using morphological variables, metabolite and phytohormone amounts, and calculating the Relative distance plasticity index (RDPI), as suggested by (Valladares, F., Sanchez-Gomez, D., & Zavala, M. A., 2006), with RDPI = ∑(dij→i’j’/(xi’j’+xij))/n), where n is the total number of distances, and j and j’ are two individuals belonging to different treatments (i and I’). This index normalizes traits with a value between 0 (no plasticity) and 1 (maximum of phenotypic plasticity) (Valladares, F., Sanchez-Gomez, D., & Zavala, M. A., 2006). The different RDPIs were calculated for morphological, metabolomic and phytohormone data by comparison of aquatic and terrestrial morphotype values in aquatic *versus* terrestrial conditions. The evolution of phenotypic plasticity between t14 and t28 was also evaluated for each data and morphotype during the time-course.

### Statistical analysis

The effect of the conditioning of both morphotypes was assessed on morphological and physiological data by running principal component analyses (PCAs) and analyses of variance (ANOVAs). ANOVA were carried out with morphotype, condition, biological replicate and morphotype x condition interaction as factors. The normality of the residuals and the homoscedasticity were verified by Shapiro–Wilk and Bartlett tests, and normality was achieved in some cases by transforming morphological traits and the metabolite concentrations. The Bonferroni test was used to compare data for different times (t14 vs t28). Statistical analyses were performed with the software R 3.5.2 with FactoMineR (Lê et al., 2008) and interface Rcmdr packages (Fox, 2005). For all ANOVA, only few biological replicate effects were observed.

For the metabolites, which varied significantly depending on the condition and morphotype, we performed pathway enrichment analyses in Metaboanalyst 3.0 (Xia et al., 2015). Fisher’s exact test algorithm was performed for these pathway analyses, with *Arabidopsis thaliana* as the reference model. In this procedure, the number of hits between the metabolites in our dataset and all metabolites of a given pathway were calculated. Metabolite data presented in the figures are untransformed. We compared: aquatic and terrestrial conditions, whatever the morphotype (Am-a and Tm-a *vs* Am-t and Tm-t), aquatic and terrestrial morphotypes, whatever the condition (Am-a and Am-t *vs* Tm-a and Tm-t), for each morphotype between condition (for example, Tm-a *vs* Tm-t) and between different morphotypes in the same condition (for example, Tm-a *vs* Am-a)

We used “ameztegui/Plasticity” packages to calculate RDPI and statistically analyses by t-test comparison between morphotypes (Am *vs* Tm) and times (t14 *vs* t28).

## Results

### Terrestrial Morphotype Exhibits Enhanced Morphological and Fitness Traits in Both Aquatic and Terrestrial Environments

Principal component analyses (PCA) were carried out and combined the measured morphological variables at days 14 and 28 (Fig. 1ab). The first axis (F1) explained 40.1% of the total variance, and separated plants grown in aquatic conditions from those grown in terrestrial conditions. Furthermore, F1 discriminated temporality for the terrestrial conditions.

Moreover, we observed that in the terrestrial conditions, morphological values of Am plants at days 14 and 28 were close to those observed at day 14 for Tm-t (Fig. 1a). The second axis explained 26.1% of the total variance and discriminated the temporality in aquatic conditions for Tm (Tm-a). Stronger variables of total biomass (TDM), number of leaves (nbL), number of buds (nbB), plant length (LP), root ratio (Rr) and root allocation (AlocR) characterised the plants growing in terrestrial conditions, while greater ratio of leaf number and of internode number to plant length (NbLLP, NbILP), shoot ratio (Sr) and shoot allocation (AlocS) characterised plants growing in aquatic conditions (Fig. 1b). Focusing on the PCA in aquatic conditions, we observed that the terrestrial morphotype had a greater number of leaves (NbL) and a higher leaf number ratio (NbLLP) at t14, whereas at t28, a higher root allocation (AlocR), plant length (LP), shoot ratio (Sr) and total dry biomass (TDM) characterised the terrestrial morphotype (Fig. S2).

**Fig. 1:**
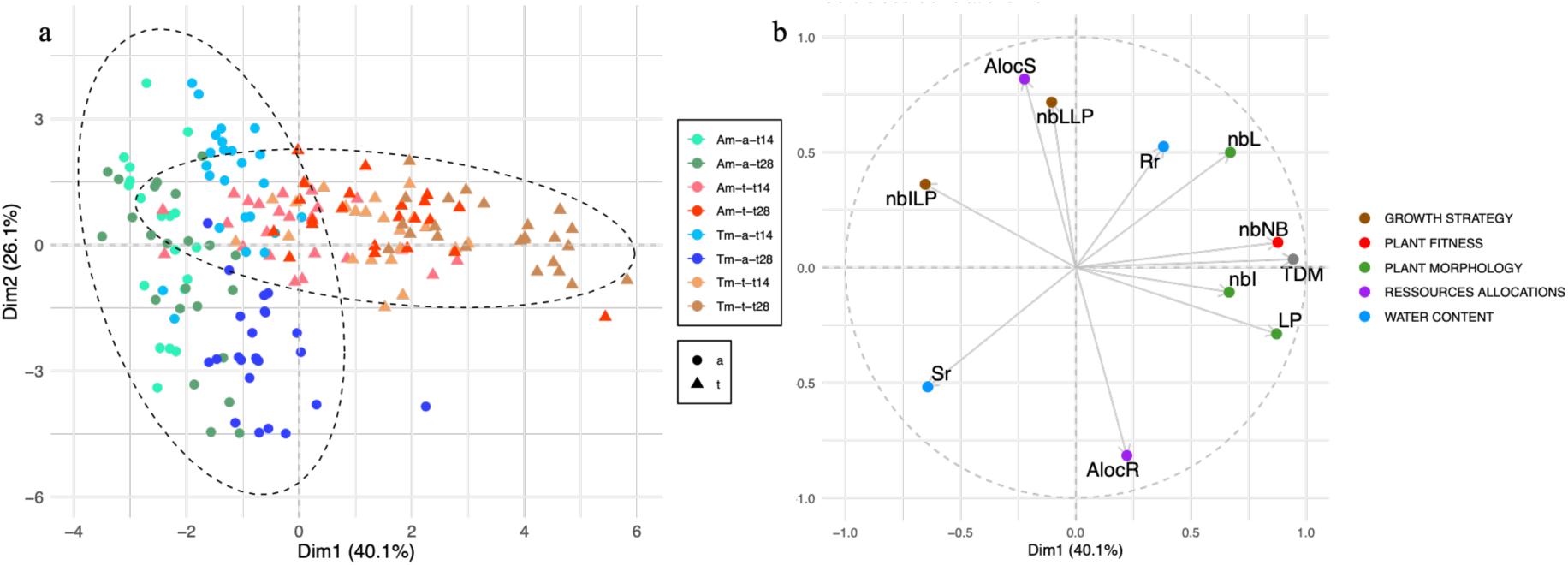
Principal component analysis (PCA) representation allowing the characterisation of the morphological variability at 14 and 28 days of experiment (-t14 and –t28) of aquatic and terrestrial morphotypes in aquatic (Am-a, Tm-a) and terrestrial (Am-t, Tm-t) conditions. (a) Individuals factor map. Each data point represented one biological repeat. (b) Variables factor map. Morphological variables represented are number of leaves (nbL), number of nodes with roots (nbNR), number of buds (nbB), fresh and dry mass of roots and shoots (FMS, FMR, DMS and DMR), shoots and roots ratio (FMS/DMS = Sr; FMR/DMR = Rr), water content (WC).

Analyses of variance were carried out for the morphotype, the condition, the time, the biological replicate and the double interactions at 14 and 28 days (Table S1; Figs. 2 and S2). We observed significant effects for at least one factor and interaction for most measured variables (p<0.05; Table S1), whereas there were few significant effects for biological replicates. Altogether, these data showed that the terrestrial morphotype presented higher values regardless of conditions (a<t) compared to the aquatic morphotype (Am<Tm). This was true for most variables attributed to the morphological and plant fitness traits, most of which increased over the duration of the experiment (t14<t28) (Table S1, Figs. 2 and S2). The only variables in favour of Am were the length of internodes (NbILp) and the root water content (Sr), especially in aquatic conditions (Fig. 2). The *Lgh* allocation resources evolved significantly differently over time, with an increasing root allocation (AlocR) and decreasing shoot allocation (AlocS), especially for Tm-a (Table S1, Figs. 2 and S3). Concerning the morphotype x condition interaction, Tm-t showed a significantly greater value for the number of buds (NbB) and for the plant fitness trait (DMT) than Am-t, Tm-a and Am-a (Table S1, Figs. 2 and S3).

**Fig. 2:**
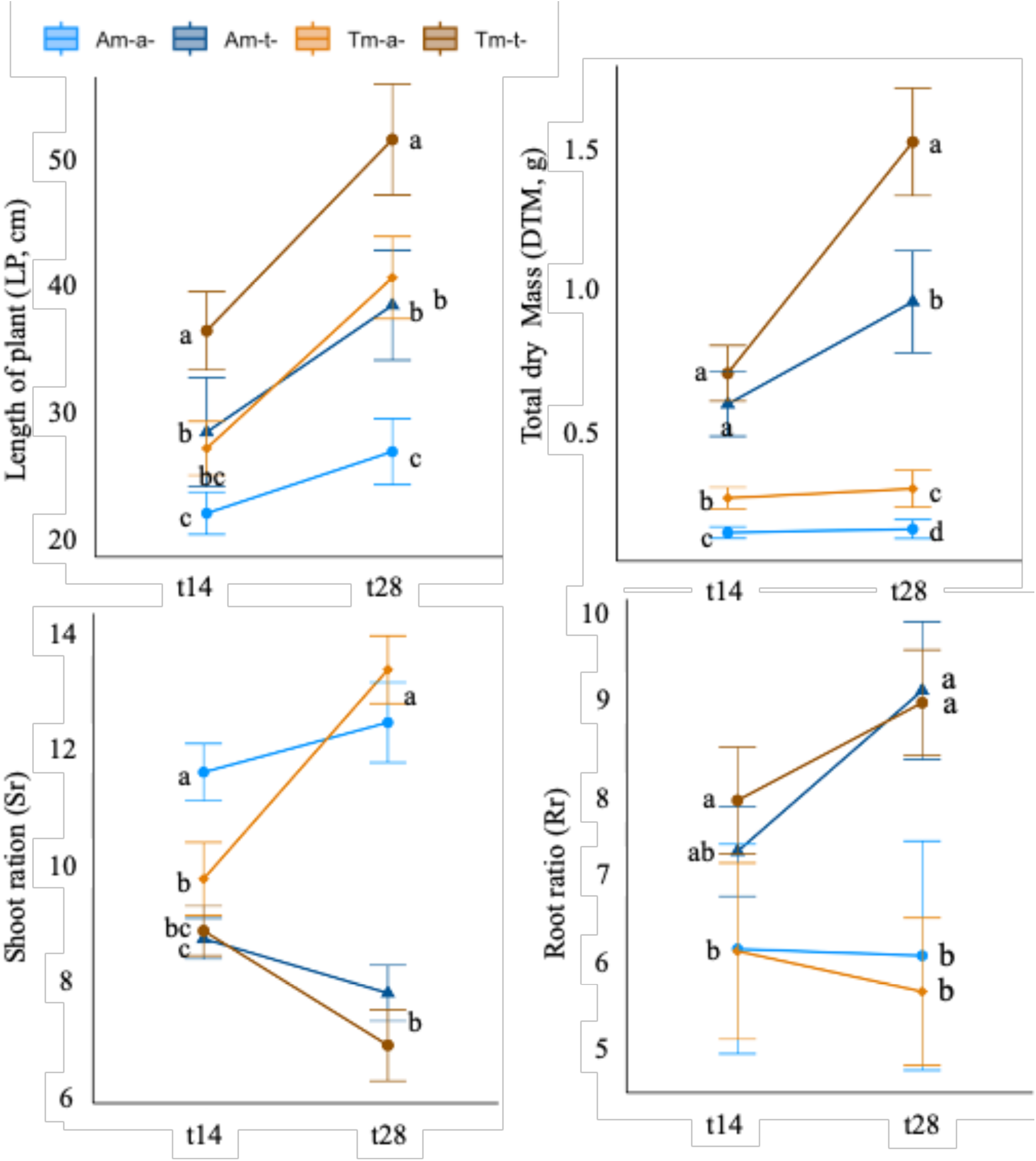
Morphological variables measured for aquatic and terrestrial morphotypes (Am: circle and Tm: triangle) in both aquatic and terrestrial conditions (-a: full line and –t: dotted line) at 14 (t14) and 28 (t28) days. Point represents the mean value of ten biological replicates with their standard errors. Letters correspond to means comparison by Tukey-test after ANOVA at t14 or t28 between morphotype*condition. Same letters mean no significant difference.

### Aquatic and Terrestrial Conditions Shape the Metabolic Responses of *Ludwigia grandiflora* subsp. *hexapetala*, with Morphotype Responses Converging Over Time

Most of the differences between morphotypes, conditions, and morphotype x condition interaction were found after 14 days of experiment (Table S2). At t28, significant morphotype and condition differences were observed, and no significant morphotype x condition interaction differences were revealed (Table S3). No significant biological replicate effect at either time was noted. Significant morphotype and/or condition effects were detected for 35 metabolites, revealing metabolite variations at t14 and t28 (Tables S2 and S4).

Principal component analyses (PCA) were carried out by combining metabolomic data at days 14 and 28 (Fig. 3). The F1 and F2 axis explained 40.1% and 26.1% of the total variance for separated aquatic and terrestrial conditions (Fig. 3a). The analysis of the correlation circle revealed that Am-t-t14 ant Tm-a mobilised particularly sugars and polyols, and amino acids and organic acids, respectively (Fig. 3b).

**Fig. 3:**
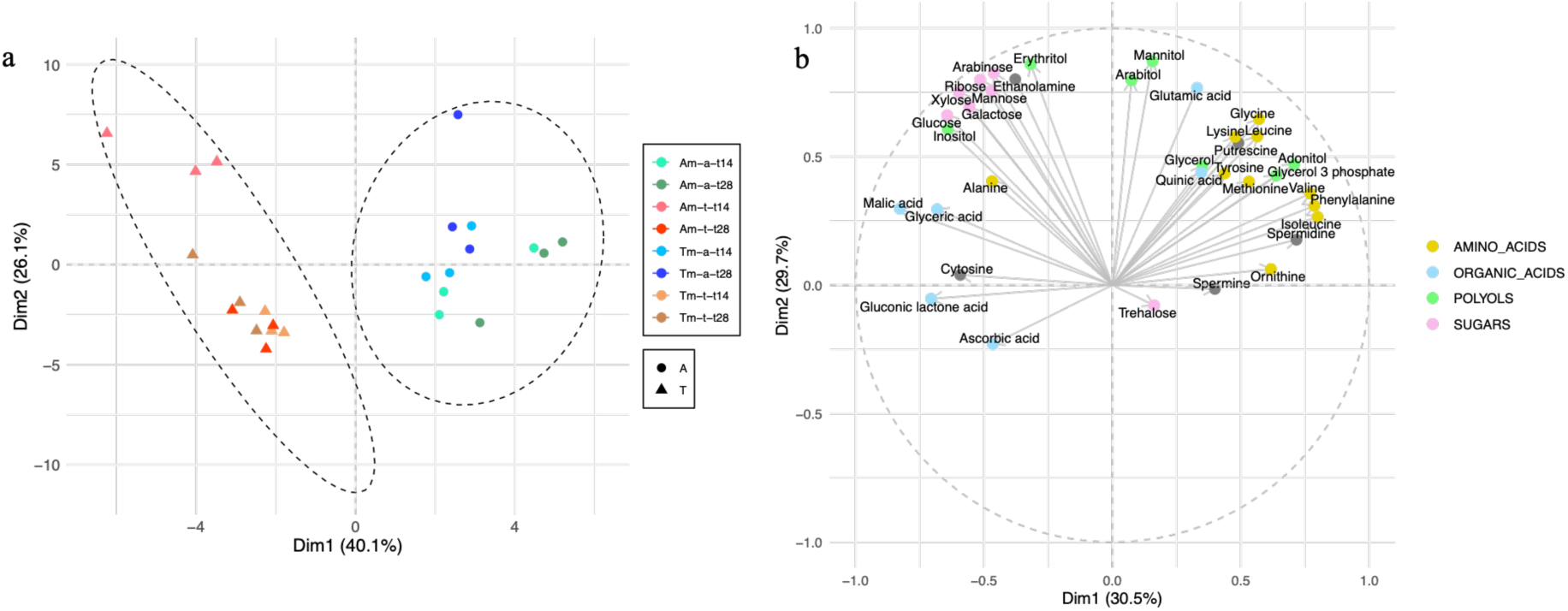
Metabolites measured in shoot of aquatic and terrestrial morphotypes (Am: circle and Tm: triangle) in both aquatic and terrestrial conditions (-a: full line and –t: dotted line) at 14 and 28 days of experiment. Metabolites showed are valine, spermidine, malic acid and mannose and their quantities were expressed in nmol/mg of dry mass. Point represents the mean value of three biological replicates with their standard errors. Letters correspond to means comparison by *Tukey-test after ANOVA* at one time between morphotype*condition. Same letters mean no significant difference.

Analyses of variance revealed that in aquatic conditions, both morphotypes (Am-a and Tm-a) had similar metabolomic responses. The plants had significantly more amino acids (glycine, isoleucine, leucine, lysine, methionine, ornithine, phenylalanine, tyrosine, valine) and polyamines (spermidine and spermine) at t14 or/and t28 compared to plants grown in terrestrial conditions (Figs. 4 and S3; Tables S2 and S3). In terrestrial conditions at t14, Am-t were characterised by significantly higher amounts of sugars arabinose, galactose, glucose, mannose, ribose, and xylose in comparison with aquatic condition (Figs. 4 and S3; Table S2). These results were consistent with those of the PCA.

**Fig. 4:**
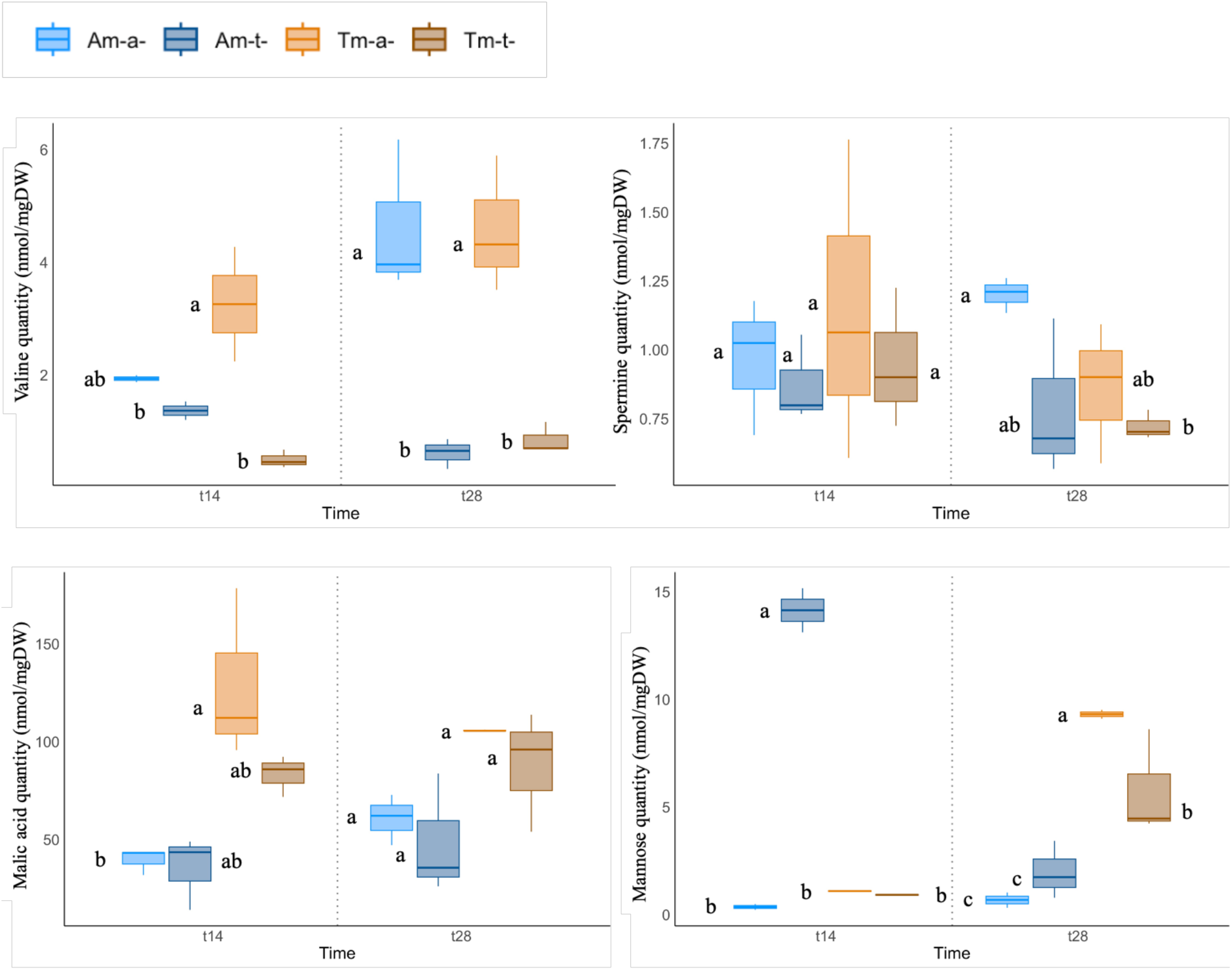
Principal component analysis (PCAs) representation allowing the characterisation of the metabolites variability at 14 (**a**) and 28 (**b**) days of experiment (-t14 and –t28) of aquatic and terrestrial morphotypes in aquatic (Am-a, Tm-a) and terrestrial (Am-t, Tm-t) conditions.

Using metabolites that discriminated the experimental conditions, we used Metaboanalyst 3.0 to describe the metabolic pathways at play at 14 and 28 days. At both times, we found four significant metabolic pathways related to the condition (a *vs* t comparison) (holm p<0.05) (Table S4). All pathways involved amino acids (Tables S2, S3 and S4, Fig. S4). In addition, at day 28, Metaboanalyst identified three specific pathways involving the phenylalanine metabolite (holm p<0.05) (Table S4, Fig. S4). These different metabolomic pathways characterised the metabolism of *Lgh* in aquatic conditions.

The terrestrial morphotype grown in aquatic conditions (Tm-a), and the aquatic morphotype grown in terrestrial conditions (Am-t) were characterised by significantly distinct metabolomic responses (Fig. 3; Fig. S4). By considering metabolites that discriminate Tm-a and Tm-t (Tm-a *vs* Tm-t and Tm-a *vs* Am-a comparison), we found that the galactose metabolism pathway was elicited. This pathway was related to Tm-a (holm p< 0.05) at 14 days corresponding to amounts of sorbitol significantly greater in Tm-a than in Tm-t and Am-a, and of glycerol 3 phosphate for Tm-t, as reported above (Fig. S4). At 14 days, the comparison of metabolites produced by Am-t and Tm-t by Metaboanalyst revealed the involvement of the glycerophospholipid metabolism pathway with ethanolamine amounts in favour of Am-t (holm p<0.05) (Fig. S3).

Both individual analysis of metabolites and metabolite pathway analysis revealed that the change observed at t14 was greater than that observed at t28 for both morphotypes in terrestrial and aquatic conditions. In fact, Am-t and Tm-t, and Am-a and Tm-a showed a closer metabolomic pattern at t28 than at t14 (Fig. 3, Tables S2 and S3).

Analyses of variances were carried out for abscisic acid and salicylic acid concentrations (Table S3). Phytohormone quantification at t28 showed a significant difference between conditions (p<0.05) for abscisic and salicylic acids, and in higher amounts in terrestrial conditions (Fig. 5, Table S3). Furthermore, we observed an interaction between morphotype x condition for auxin quantity, with significant differences between aquatic morphotypes in both conditions, in favour of aquatic conditions (Am-a *vs* Am-t) (Fig. 5; Table S3). No significant differences were observed for biological replicates.

**Fig. 5:**
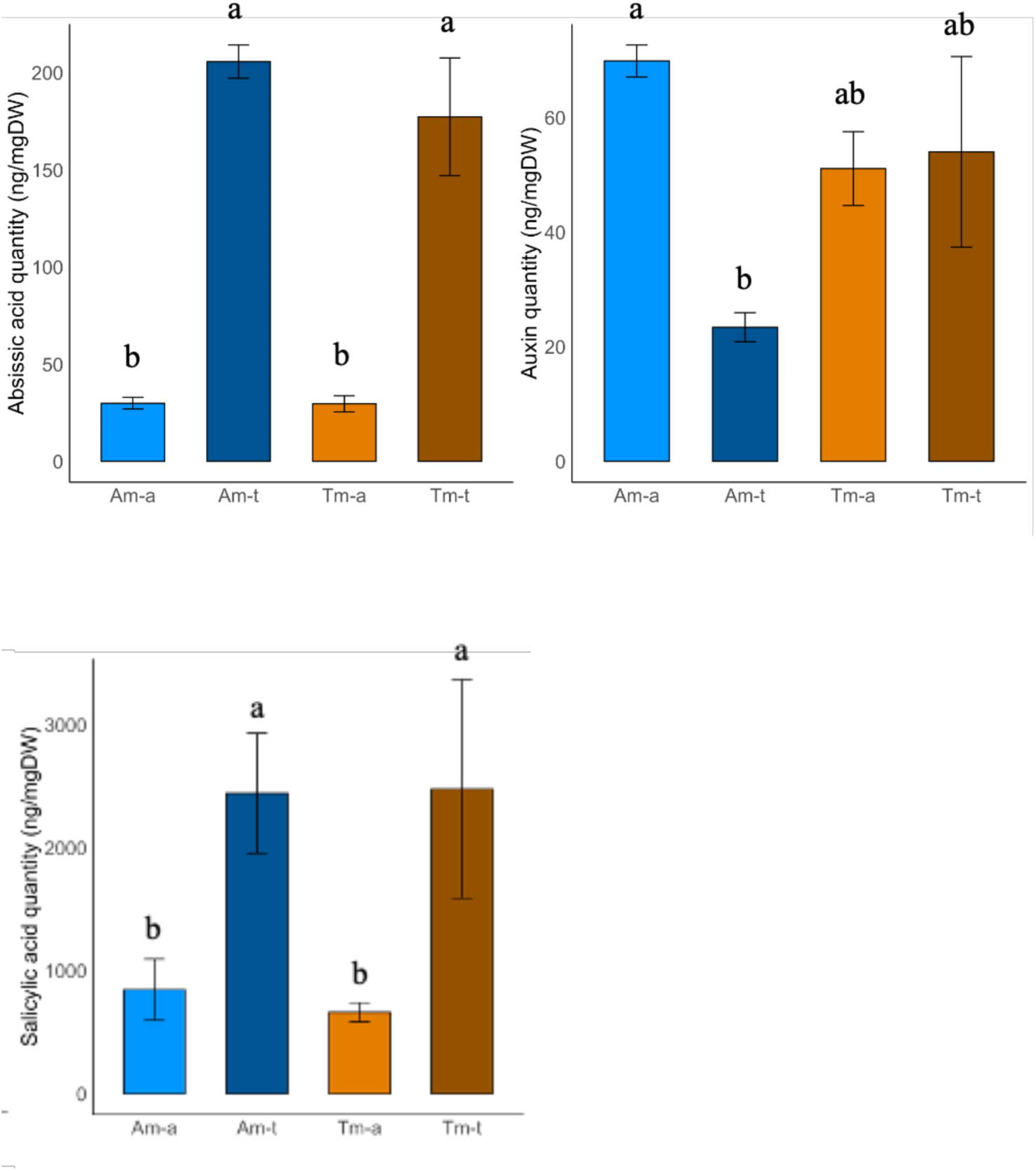
Phytohormones quantified in shoot of aquatic and terrestrial morphotypes (Am and Tm) in both aquatic and terrestrial conditions (-a and -t) at 28 days. Phytohormones quantities were expressed in ng/g DW of dry mass (DW). Point represents the mean value of three biological replicates with their standard errors. Letters correspond to means comparison by Tukey-test after ANOVA at t28 between morphotype*condition. Same letters mean no significant difference (p-value≤0.05).

### The Aquatic Morphotype Exhibits Greater Phenotypic Plasticity Than the Terrestrial Morphotype, with Plasticity Increasing Over Time

The phenotypic plasticity for all morphological variables studied was calculated using the Relative distance plasticity index (RDPI), and a *t*-test for statistical analyses. We observed significant differences in RDPI between Am and Tm, most in favour of Am, for 6 or 8 of the 11 variables at 14 and 28 days, respectively (p<0.05; Table 1). RDPI values varied between 0 and 1, with the morphological variable highly plastic (RDPI> 0,9) for number of buds, moderately plastic (0.4>RDPI>0.8) for plant fitness (total dry biomass variable) and number of leaves for Tm at t28 only, and lower plastic (RDPI<0.4) for the three morphological variables (LP, NbI, NbLLP), the water content variable (Sr) and allocation resources (AlocR, AlocS) (Table 1). No significant RDPI difference was noted between morphotypes for the ratio of internodes/plant length (nbILP) or the water content ‘root ratio’ (Rr) at either time. Thus, the morphological trait ‘number of buds’ (nbNB), the plant fitness (TDM) and the allocation resources variables (Aloc S, AlocR) showed a greater phenotypic plasticity for the aquatic morphotype at both times (Table 1, Fig. 2). For the number of leaves, Am showed a higher RDPI value at t14, whereas at t28, the Tm value was higher. Similarly, the RDPI of water content in shoots was higher at t14 for Am and then higher at t28 for tm.

**Table 1.**
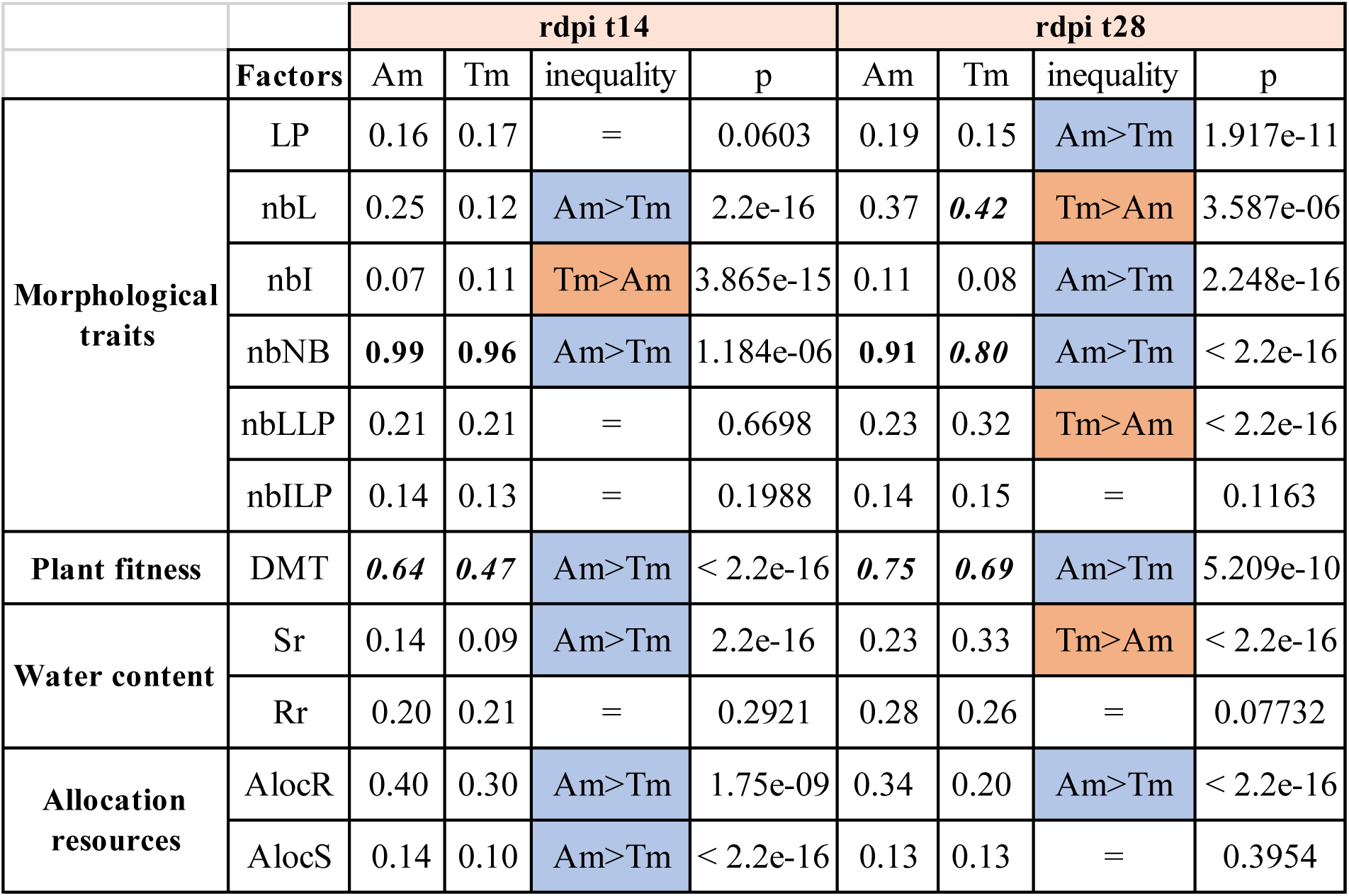
RDPI results and comparison of traits variables of both morphotypes (Am and Tm) at 14-28 days. Significance codes: .001 “***”; .01 “**”; .05 “*”; nonsignificant “=” t-test.

Comparison of RDPI at 14 and 28 days was carried out to analyse the phenotypic plasticity evolution over time (*t*-test). We noted a significant increase in RDPI between 14 and 28 days for most of morphological variables and for both morphotypes, except for the number of buds (nbNB) and the root allocation resources (AlocR) (p<0.05; Table 2). The terrestrial morphotype showed a low plasticity at t28 for the length of the plant (LP) and the number of internodes (nbI).

**Table 2:** RDPI dynamics during experimental time of traits variables of both aquatic and terrestrial morphotypes (Am and Tm, respectively). Significance codes: .001 “***”; .01 “**”; .05 “*”; nonsignificant “=” t-test.

|  |  | rdpi dynamics Am |  |  |  | rdpi dynamics tm |  |  |  |
| --- | --- | --- | --- | --- | --- | --- | --- | --- | --- |
|  | Factors | t14 | t28 | inequality | P | t14 | t28 | inequality | p |
| **Morphological traits** | LP | 0.16 | 0.19 | t28>t14 | <1.1e-3 | 0.17 | 0.15 | t14>t28 | <1.0e-4 |
|  | nbL | 0.25 | 0.37 | t28>t14 | < 2.2e-16 | 0.12 | **0.42** | t28>t14 | < 2.2e-16 |
|  | nbI | 0.07 | 0.11 | t28>t14 | 3.3e-14 | 0.11 | 0.08 | t14>t28 | < 2.2e-16 |
|  | nbNB | **0.99** | **0.91** | t14>t28 | 1.1e-3 | **0.96** | **0.80** | t14>t28 | < 2.2e-16 |
|  | nbLLP | 0.21 | 0.23 | t28>t14 | 6.6e-3 | 0.21 | 0.32 | t28>t14 | <1.6e-45 |
|  | nbILP | 0.14 | 0.14 | = | 0.6156 | 0.13 | 0.15 | t28>t14 | <5.6e-4 |
| **Plant fitness** | DMT | **0.64** | **0.75** | t28>t14 | <1.7e-18 | **0.47** | **0.69** | t28>t14 | <3.3e-74 |
| **Water content** | Sr | 0.14 | 0.23 | t28>t14 | < 2.2e-16 | 0.09 | 0.33 | t28>t14 | < 2.2e-16 |
|  | Rr | 0.20 | 0.28 | t28>t14 | <1.5e-14 | 0.21 | 0.26 | t28>t14 | <6.0e-9 |
| **Allocation resources** | AlocR | 0.40 | 0.34 | t14>t28 | <4.5e-4 | 0.30 | 0.20 | t14>t28 | <1.5e-22 |
|  | AlocS | 0.14 | 0.13 | = | 0.4463 | 0.10 | 0.13 | t28>t14 | <1.9e-08 |

A similar approach was used for phytohormone and metabolite RDPI evaluations. Among the 59 metabolites analysed, 36 showed at least one significant difference for RDPI values at t14, t28 or over the time of the experiment (p<0.05; Tables S5 and S6). We observed significant differences in RDPI for 25, and only ten of the 59 metabolites analysed at 14 and 28 days, respectively (p<0.05; Table S5). RDPI values varied between 0 and 1, with highly plastic for sugars particularly for Am at 14 days (RDPI> 0,9), moderately plastic for a majority of metabolites at 14 days, especially for amino acids and organic acids (0.4>RDPI>0.8), and lower plastic for some metabolites such as spermidine and spermine (RDPI<0.4). As we only had phytohormone assays at t28, the RDPI was calculated for this time. Only auxin showed a significant difference, with a moderate plasticity in favour of Am.

The number of metabolites with significantly greater phenotypic plasticity was higher for the aquatic morphotype than for the terrestrial morphotype at both times, with the RDPI of 17 metabolites superior in Am compared to eight in Tm at t14. At t28, the RDPI of six metabolites and auxin were superior in Am compared to three in Tm. Thus, at 14 days, the majority of sugars and organic acids showed a higher RDPI for the aquatic morphotype compared to the terrestrial one (p<0.05; Table S5; Fig 3). A greater plasticity for the terrestrial morphotype was observed for only eight metabolites (three amino acids; two polyols, two organic acids and putrescine) (p<0.05; Table S5; Fig. S4).

To assess the dynamics of plasticity expression for each metabolite over time, we compared RDPI values at t14 and t28 for Am and Tm, respectively. We observed a significantly greater amino acid RDPI at t28 than at t14 for both morphotypes, with higher and moderate plasticity, in particular for glycine, methionine, and phenylalanine (p<0.05; Table S6; Fig. S4). For the majority of sugars, however, the RDPI between 14 and 28 days decreased significantly in the aquatic morphotype (t14>t28) and increased significantly in the terrestrial morphotype (t28>t14) (p<0.05; Table S6; Fig. S4).

All these data showed that Am presented a higher phenotypic plasticity for both morphological traits and metabolites compared to Tm, especially at t14. However, phenotypic plasticity of both morphotypes increased similarly over time for both traits and metabolites, except for some sugars and organic acids in favour of Tm.

## Discussion

To explore the acclimatation responses of *Lgh* when grown in terrestrial conditions, we measured morphological and metabolomic responses of *Lgh* in both habitats (aquatic and terrestrial), and evaluated the phenotypic and metabolomic plasticity of both morphotypes in their original or reverse habitats.

### *Ludwigia grandiflora* subsp. *hexapetala* Appeared to be Pre-adapted to the Terrestrial Habitat and Able to Adjust its Growth and Metabolism to its Environment

To survive drought events, several aquatic plants reduce leaf area, increase relative root biomass (root/shoot ratio) and reduce transpiration rates to increase water use efficiency (Hussner et al., 2009; N. Venter, B.W. Cowie, E.T.F. Witkowski, G.C. Snow, M.J. Byrne, 2017; Touchette et al., 2007). For instance, in the two aquatic invasive macrophytes, the water hyacinth, *Nymphoides peltata,* and *Myriophyllum aquaticum*, similar morphological changes were observed in function of water availability. In terrestrial habitats, *N. peltata* showed lower total biomass, higher root biomass location, and root-shoot ratio, than in aquatic habitats (Z. Li et al., 2011; Venter et al., 2017). Similarly, the amphibious plant, *M. aquaticum* accumulated less biomass when water levels declined (Hussner et al., 2009). In our study, a distinct pattern was observed for *Lgh,* which produced a higher total biomass in terrestrial habitats than in aquatic habitats. One explanation could be that the *Lgh*’s ancestral terrestrial origin, reported by Bedoya et al (2015), could favour its development in terrestrial habitats. The phylogenetic construction of *Lgh*, according to its morpho-anatomical character, suggests various back and forth between the terrestrial and aquatic environments, which could explain the pre-adaptation enabling it to colonise terrestrial environments (Bedoya & Madriñán, 2015). Another argument for the pre-adaptation of *Lgh* is the fact that no significant difference was observed for resource allocations between morphotypes, indicating that *Lgh* is able to allocate similar resources to different parts of the plant independently of the environment.

The ability of *Lgh* to adapt to a wide range of habitats is likely due to the physiological plasticity of the plant. Our metabolomic analyses were in line with those obtained by Billet et al, (2018) and evidenced that the transition of *Lgh* from aquatic habitats to terrestrial habitats is supported by metabolic adjustments. In aquatic conditions, a mobilisation of the pathway involving amino acids, in particular phenylalanine, and higher quantities of auxin and polyamines were highlighted. Amino acids are associated to low concentration of C0_2_ and tolerance to hypoxia, as is the case in aquatic habitats (Das & Uchimiya, 2002; Hussner, Mettler-Altmann, Weber, & Sand-Jensen, 2016; Less & Galili, 2008; Limami, 2014; Pedersen & Colmer, 2014). Furthermore, phenylalanine, tryptophane and tyrosine are involved in protein synthesis, and are precursors for various plant hormones such as auxin, which play an important role in flooding (Sasidharan *et al*., 2018; Fukao *et al*., 2019). In addition, polyamine production is important in many processes, such as stress responses, morphogenesis, and growth (Kusano et al., 2008; Vera-Sirera et al., 2010). In terrestrial conditions, a wide range of sugars and organic acids, and higher quantities of abscisic acid and salicylic acid were observed. Abscisic acid and salicylic acid are both involved in drought stress and developmental processes, such as stomatal closure or heterophylly (Mommer & Visser, 2005; Prodhan et al., 2018). Thus, in *Ludwigia arcuate*, the formation of terrestrial leaves on submerged shoots was induced by ABA (Kuwabara et al., 2003). In addition, Sami *et al*. (2016) showed the crosstalk between phytohormones such as ABA, and sugar metabolites in drought stress.

All our results suggested that *Lgh* grown in aquatic or terrestrial habitats can develop many strategies in terms of morphological and metabolomic traits in order to adapt to stressful environments. These results also support our hypothesis that *Lgh* exhibits high phenotypic flexibility according to its habitat, and the terrestrial morphotype presented higher trait values than in aquatic conditions.

### The Reversibility of *Lgh*’s Plastic Response to Changing Conditions Depends on Morphotypes

The terrestrial morphotype exhibits greater/ higher, morphological trait values than the aquatic morphotype in aquatic conditions, while the aquatic morphotype in terrestrial environments shows lower morphological trait values than those observed for the terrestrial morphotype. In addition, in aquatic conditions, the terrestrial morphotype showed mixed physiological responses between Tm-t and Am-a. On the contrary, when the aquatic morphotype in terrestrial conditions mobilised specific metabolomic pathways, this did not seem to have a positive effect on the growth and development of the aquatic morphotype. These observations questioned the reversibility of *Lgh*’s plastic response to changing conditions and suggested two possible explanations.

Firstly, this may be the environmental conditions, which, as a constraint, lead the population to acclimatize to its environment through the expression of phenotypic plasticity before natural selection and hence local adaptation. Secondly, although numerous studies of flooding responses have been carried out, little is known about the recovery process during the post flood period (Yeung *et al*., 2018; Yeung, Bailey-Serres, & Sasidharan, 2019). Hilker & Schmülling (2019); Maury *et al*. (2019); Mozgova, Mikulski, Pecinka, and Farrona (2019) suggested the possibility that plants maintain a memory of the constraint (memory stress or priming). During its life cycle, the *Lgh* terrestrial morphotype undergoes two predictable flooding phases: the submergence phase, and the post-submergence recovery phase. The latter may be more stressful than the submergence phase for *Lgh*, and may also induce memory stress, conditioning its ability to grow in aquatic and terrestrial environments.

Contrary to our third hypothesis, the terrestrial morphotype did not completely converge to the aquatic morphotype in the reverse condition, especially in terms of adjusting its metabolism.

#### Both Phenotypic Plasticity and Local Adaptation Contribute to Invasiveness of *Lgh*

Rapid adaptive evolution and phenotypic plasticity were recognized as two mechanisms involved in the invasiveness of invasive plants (Xiong et al., 2024). Szabó *et al*. (2019) studied two macrophyte invasive species: *Elodea nuttallii* and *Elodea canadensis*. They showed that *Elodea nuttallii’*s phenotypic plasticity is greater than that of *Elodea canadensis,* which in turn contributes to its invasive success. Furthermore,(Xiong et al., 2024) demonstrated that rapid adaptive evolution and phenotypic plasticity contribute to the invasiveness of *Ambrosia artemisiifolia* under different levels of N availability, suggesting that these two factors can enable invasive plants to colonise a wide range of environmental conditions.

In this study, the first to report on *Lgh*’s phenotypic plasticity, revealed that phenotypic plasticity was expressed when the environment changed, and in function of the morphotype. This reinforces our first hypothesis. However, the aquatic morphotype of *Lgh* showed a greater phenotypic plasticity than the terrestrial morphotype for the majority of morphological variables, metabolites amounts and auxin quantities. This result invalidated our second hypothesis (2), according to which the plasticity of an organism developing in heterogeneous conditions is greater than that of an organism in a homogenous environment, which is the case for the terrestrial morphotype, as it is subject to water level seasonality (Murren et al., 2015). But according to Matzek (2012) it is trait values and not trait plasticity which explain the performance of invasive species in changing environments that was observed for the terrestrial morphotype in our study.

In this study, we postulated that phenotypic plasticity explains the rapid colonisation of terrestrial environments. However, local adaptation is also considered to be an important mechanism in the invasiveness of a species. (Savić et al., 2024) observed, after a 10-year residence time of *Ambrosia trifida* populations in the invasion area, a reduction in phenotypic plasticity associated with more vigorous vegetative growth, which is similar to results obtained in our study. These authors suggested that the heritability of the favourable traits allows local adaptation.

According to Lande’s hypothesis (Lande, 2015), which suggests that phenotypic plasticity is temporally dynamic, we can imagine that the phenotypic plasticity of *Lgh* has evolved. Thus, at the beginning of terrestrial colonisation between 1995-2000 in France, the terrestrial morphotype of *Lgh* could present a strong plasticity. This phenotypic plasticity could then have evolved and integrated, which explains the lower RDPI observed for the terrestrial morphotype. On the other hand, for the aquatic morphotype, the higher phenotypic plasticity observed could reflect stress responses alongside its ability to invade new habitats. In addition, the morphological trait values of the aquatic morphotype are lower than those of the terrestrial morphotype, which may reflect the cost of plasticity (Murren et al., 2015). To artificially mimic the transition from aquatic to terrestrial morphotypes, experiments could be carried out by constraining the aquatic morphotype to terrestrial conditions and assessing the evolution of its phenotypic plasticity.

## Conclusion

In this work, we demonstrated the existence of distinct morphological and physiological traits, as well as variations in phenotypic plasticity between aquatic and terrestrial morphotypes of the aquatic invasive plant, *Ludwigia grandiflora* subsp. *hexapetala* in response to changing environments. The terrestrial morphotype showed higher trait values but lower phenotypic plasticity than the aquatic morphotype, suggesting a pre-adaptation of *Lgh* to terrestrial habitats, and involvement of phenotypic plasticity and local adaptation in its invasiveness. The ability of the terrestrial morphotype to adapt quickly to a new environment, as well as a possible aquatic return with increased capacities, is worrisome for the management of the invasion.

To carry out the phenotypic plasticity approach as suggested by Lande, (2015), a comparison of the phenotypic plasticity of several *Lgh* populations with various introductory times would make it possible to assess the invasion time. Furthermore, phenotypic plasticity is notably under genetic and epigenetic controls within this framework (Duncan et al., 2014). To disentangle the mechanisms involved in plasticity evolution during invasion, Hodgins et al., (2025) recommended examining both genomes and epigenomes. *Ludwigia grandiflora* subsp. *hexapetala* exhibits both genetic diversity and a potentially significant epigenetic effect and shows itself to be capable of rapid adaptation, is a good model for exploring the evolutionary dynamics of invasive species (Richards *et al*., 2017; Marin *et al*., 2019; Genitoni *et al*., 2020).

## Acknowledgements

The authors would like to thank Michel Bozec for his help in collecting *Lgh* plants in natura and obtaining morphological data. This research was supported by ONEMA (French National Agency to Water and Aquatic Environments). Julien Genitoni received a Ph.D. grant from INRAE Département ECODIV—Région Bretagne. The authors thank the Experimental Unit of Aquatic Ecology and Ecotoxicology (U3E) 1036, INRAE, which is part of the research infrastructure Analysis and Experimentations on Ecosystems-France, for help with the maintenance of plants. The authors thank S. Citerne for hormonal dosages (IJPB’s Plant Observatory Chemistry/Metabolism platform, INRAE Versailles).

## Author contributions

Dominique Barloy, Stéphane Maury and Julien Genitoni conceived and designed the experiment. Julien Genitoni, Daniel Vassaux and Dominique Barloy contributed to collecting morphological data and to sampling for metabolomic analysis. Julien Genitoni and David Renault carried out metabolomic analysis. All authors contributed to analysis of the data. Julien Genitoni wrote the first draft of the manuscript supervised by Dominique Barloy and Stéphane Maury. All authors contributed to revisions and approved the final manuscript.

