## Supplemental Figures S1-S4 for "Adaptive Strategies of the invasive aquatic plant, *Ludwigia grandiflora* subps. *hexapetala*: Contrasting Plasticity Between Aquatic and Terrestrial Morphotypes"

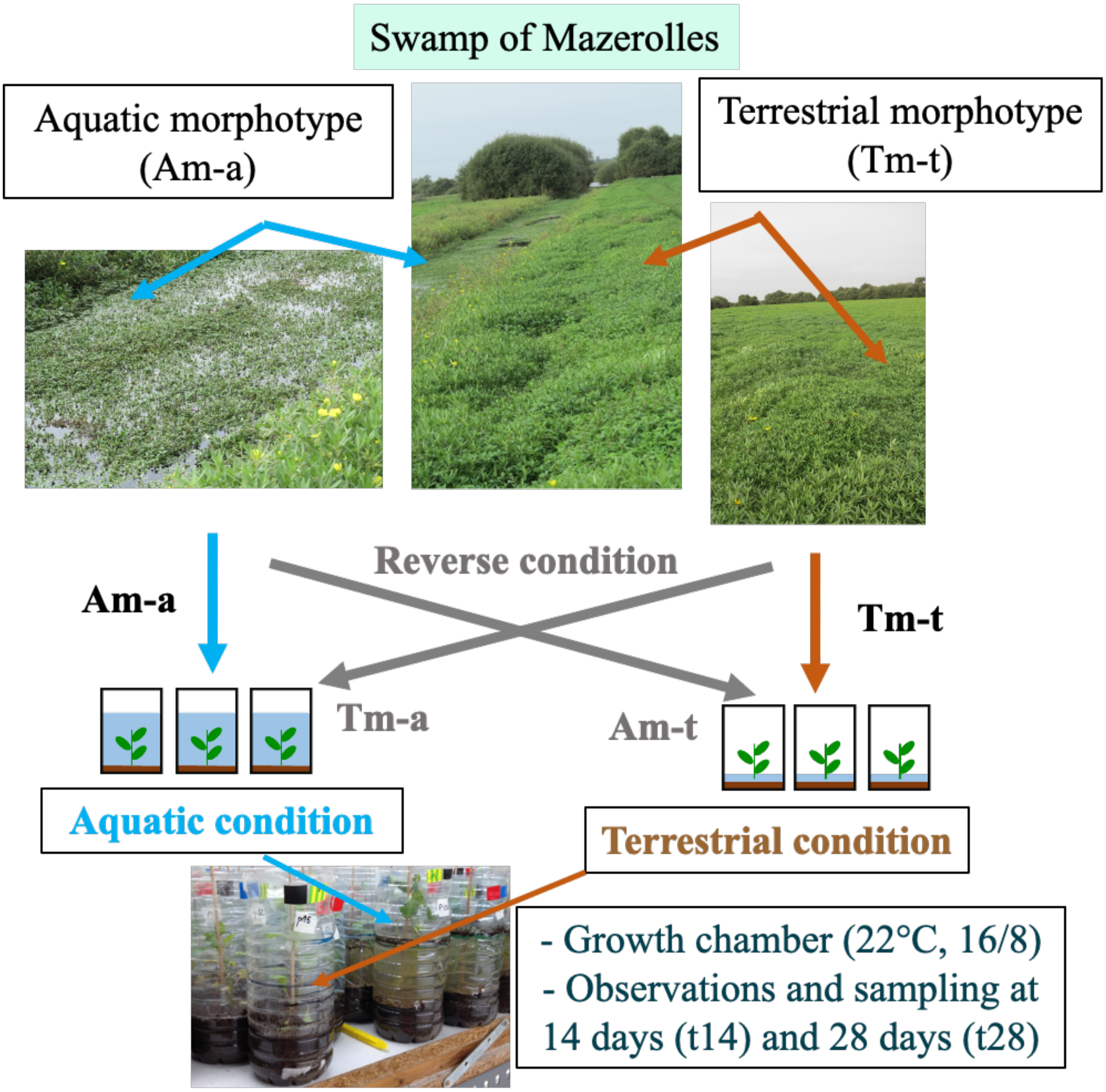


Figure S1: Experimental design

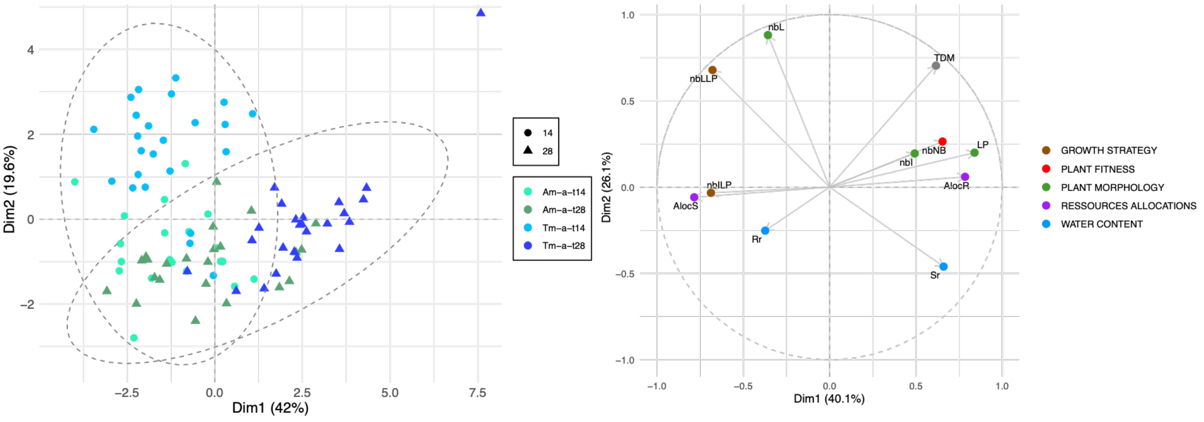


Figure S2: Principal component analysis (PCA) representation allowing the characterisation of the morphological variability at 14 and 28 days of experiment (-t14 and –t28) of aquatic and terrestrial morphotypes in aquatic (Am-a, Tm-a) condition. (a) Individuals factor map. Each data point represented one biological repeat. (b) Variables factor map. Morphological variables represented are number of leaves (nbL), number of nodes with buds (nbNB), number of internode (NbI) length of shoot (LP), shoot and root ratios (Sr = fresh mass of shoots/dry mass of shoots; Rr = fresh mass of roots/dry mass of roots), leave and Internode numbers ratios (nbLLP=nbL/LP; nbILP=nbILP), Total dry mass (TDM), shoot and root resource allocations (AlocS=dry mass of shoot/TDM; AlocR=dry mass of roots/TDM).

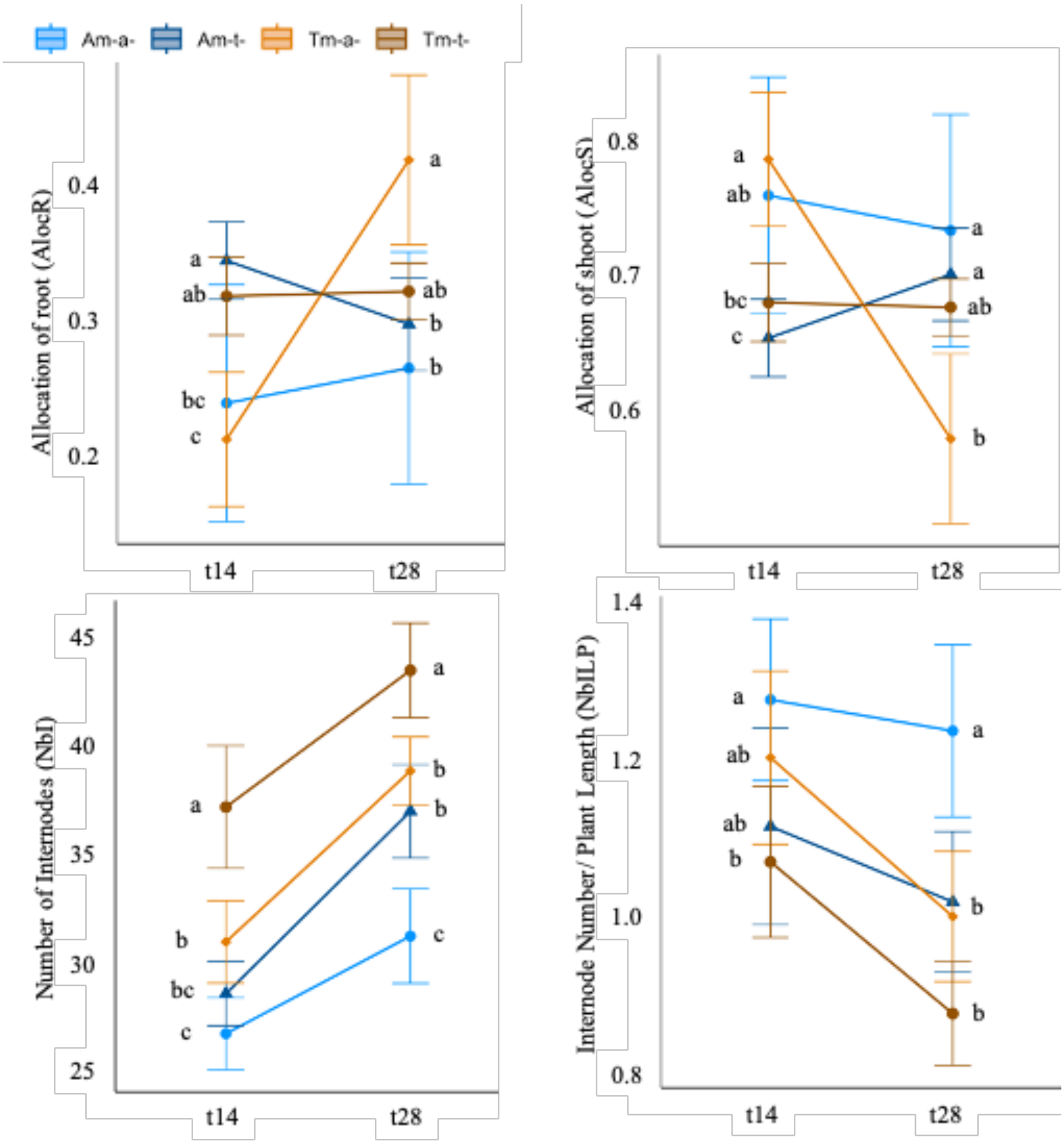


Figure S3: Morphological variables measured for aquatic and terrestrial morphotypes (Am: circle and Tm: triangle) in both aquatic and terrestrial conditions (-a: blue color and -t: orange color) at 14 (t14) and 28 (t28) days. Point represents the mean value of two biological replicates with their standard errors. Letters correspond to means comparison by Tukey-test after ANOVA at t14 and t28 between morphotype*condition. Same letters mean no significant difference (p-value≤0.05).

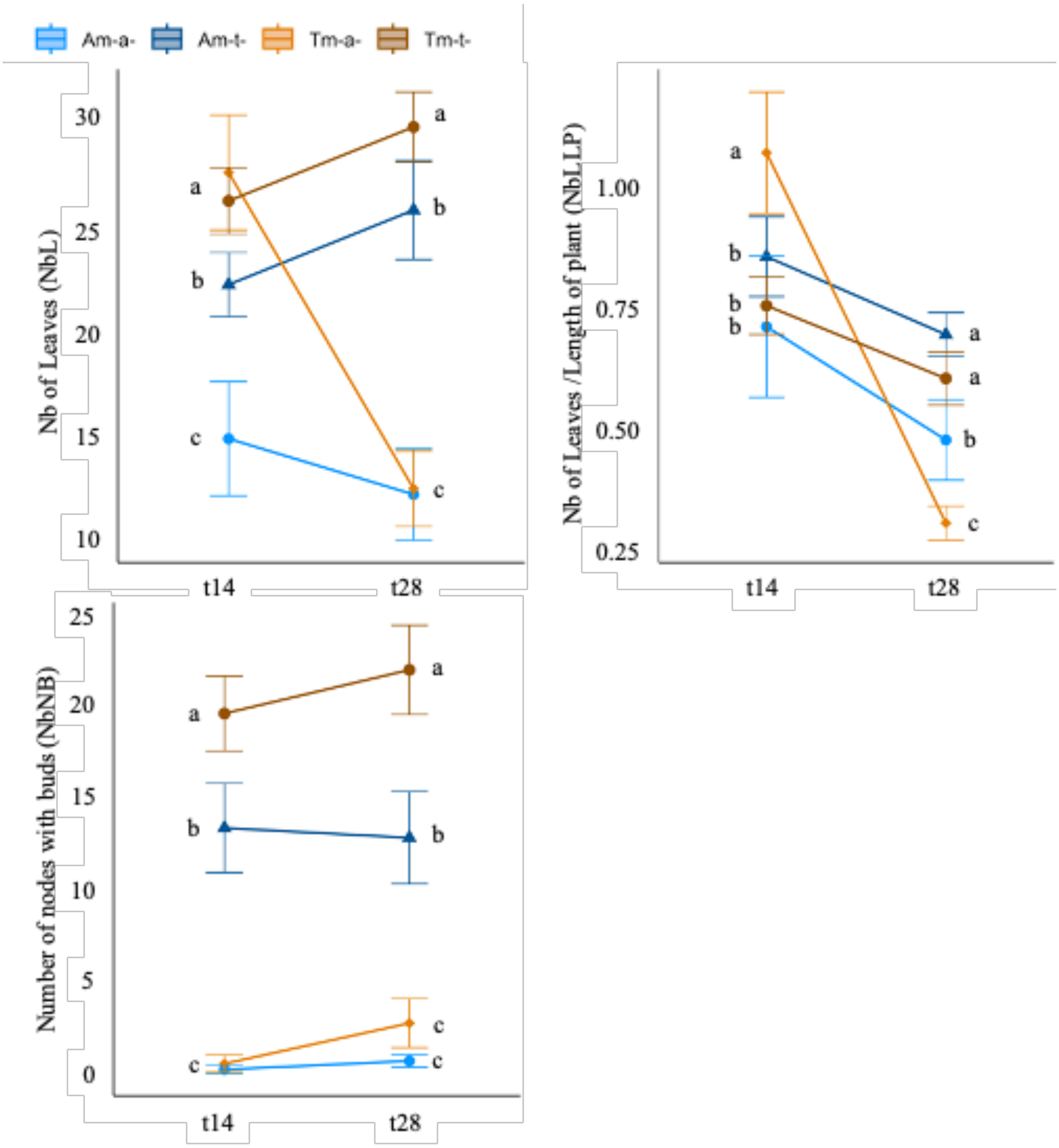


Figure S3: continued

a) Amino acids

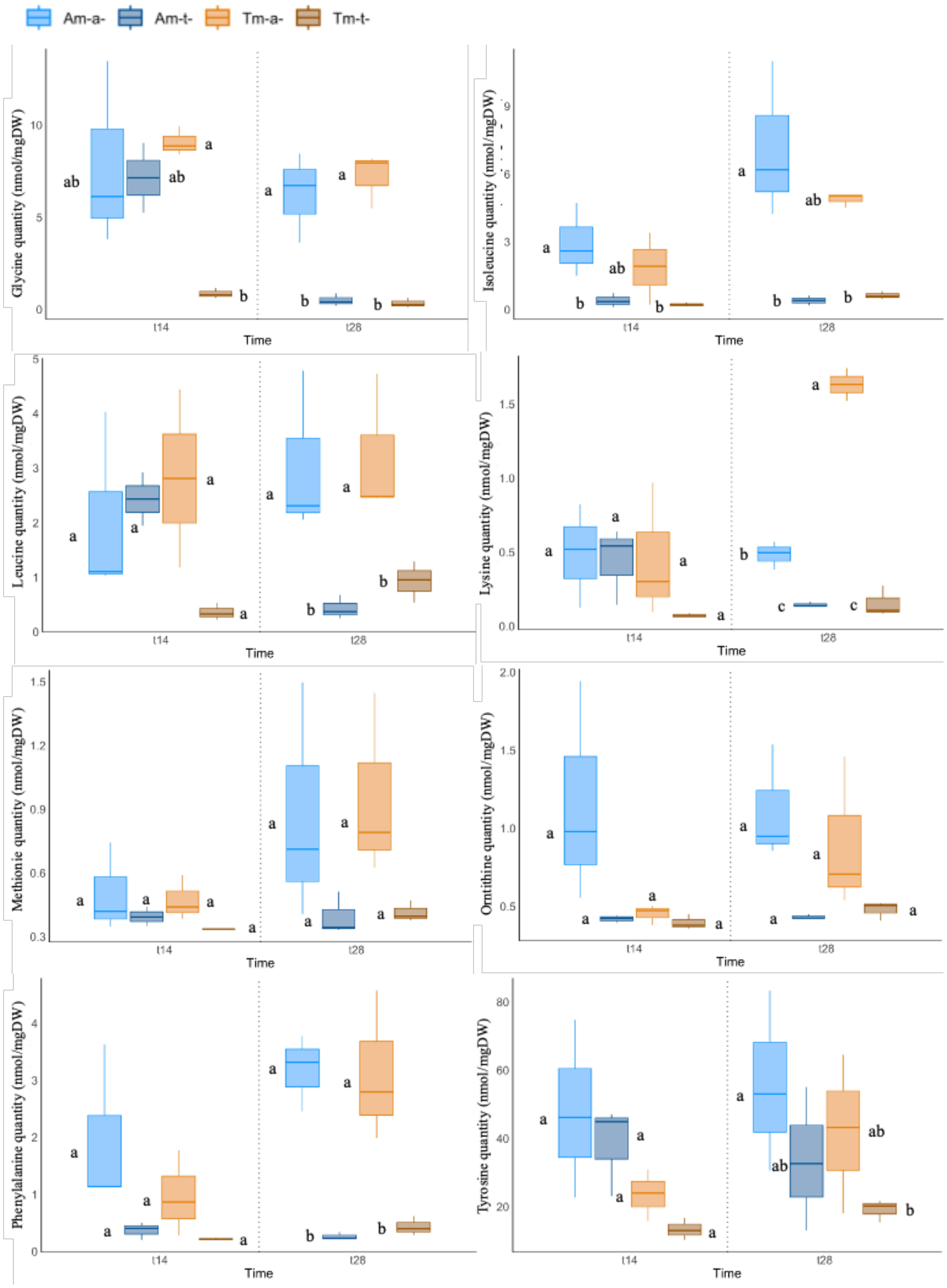

Figure S4: Metabolites measured in shoot of aquatic and terrestrial morphotypes (Am: Blue and Tm: orange) in both aquatic and terrestrial conditions (-a: light colour and -t: dark colour) at 14 (t14) and 28 (t28) days. a) amino acids.

Metabolite quantities were expressed in nmol/mg of dry mass. Box plot represents the median values of three biological replicates (horizontal line). Box extend from 25$^{th}$ to 75$^{th}$ percentile of value distribution. Vertical extending lines denote adjacent values (i.e., the most extreme values within 1.5 interquartile range of the 25$^{th}$ and 75$^{th}$ percentile of each value). Letters correspond to means comparison by Tukey-test after ANOVA at t14 and at t28 between morphotype*condition. Same letters mean no significant difference ($p$-value$\leq$0.05).

b) Polyols

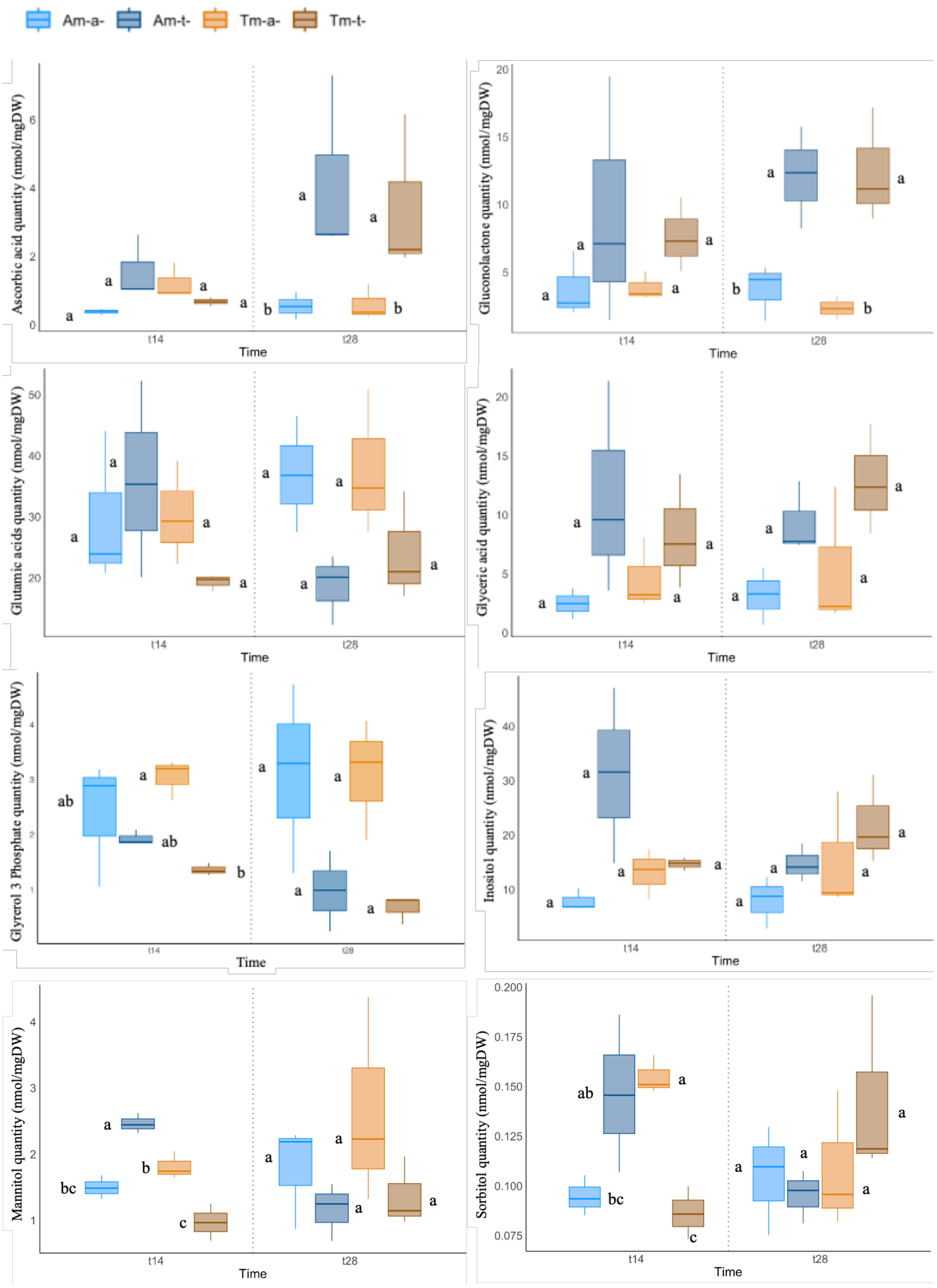


Figure S4: continued; b) Polyols

c) Organic acids

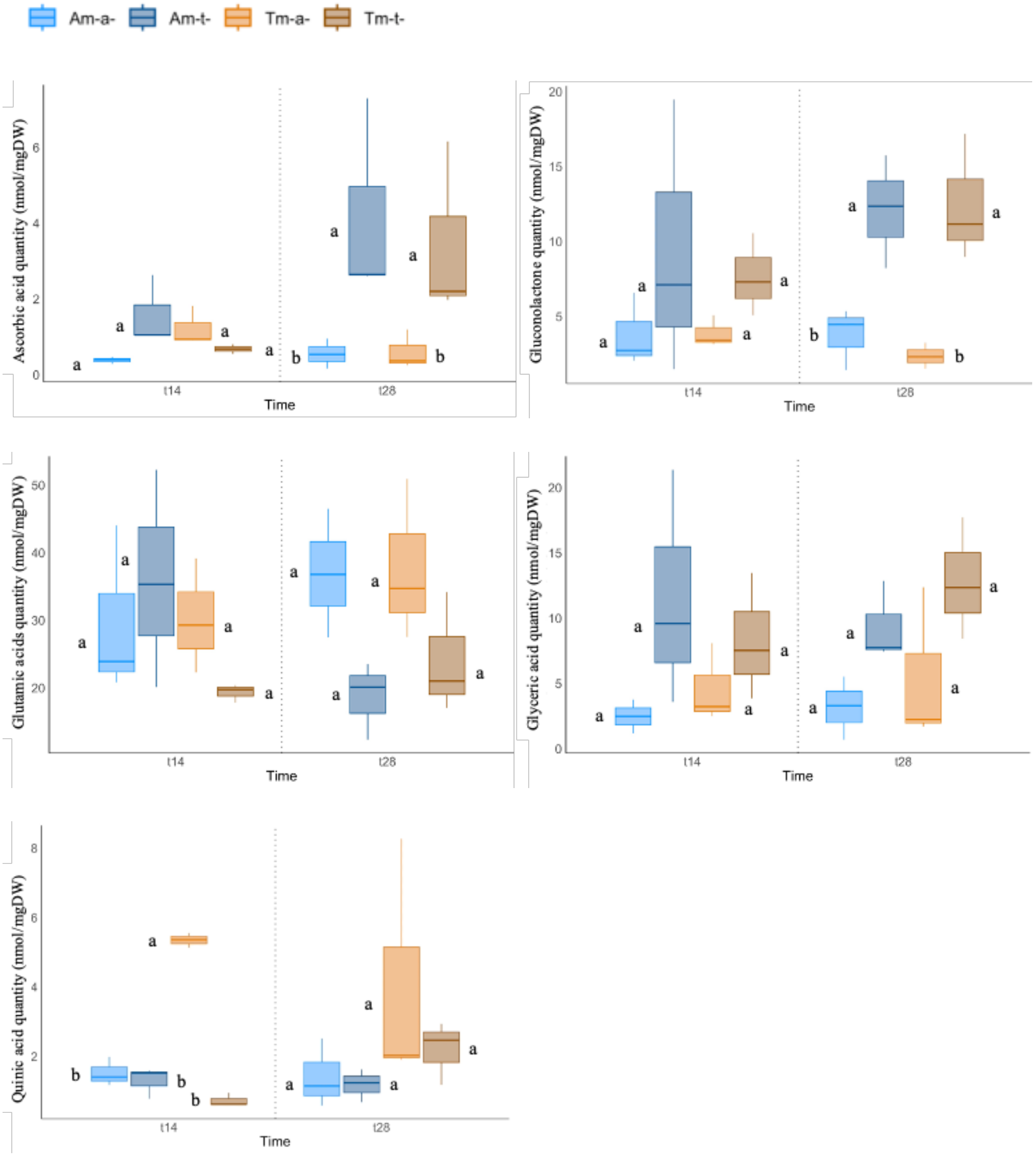


Figure S4: (c) Organic acids

d) sugars

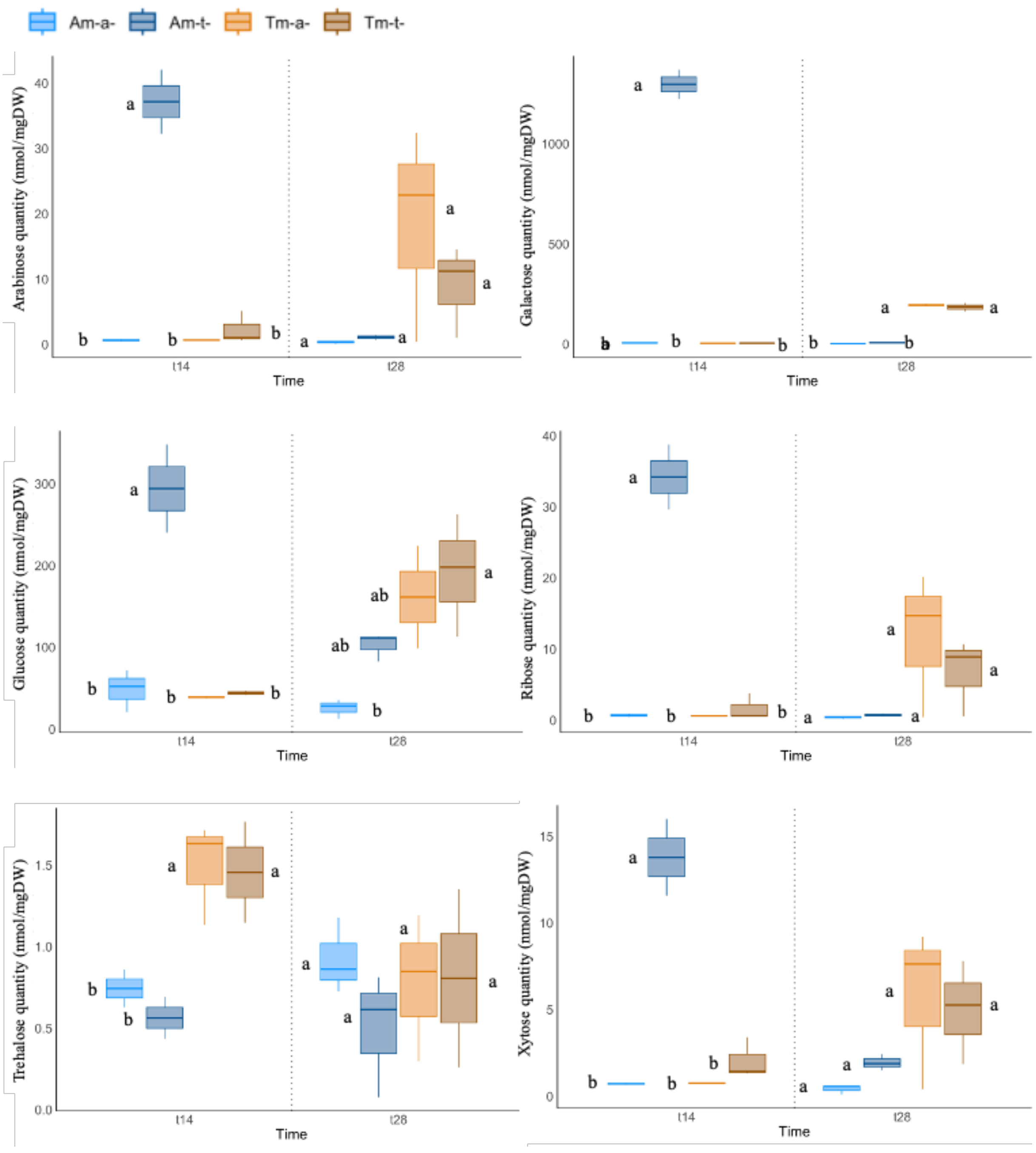


Figure S4: continued, d) Sugars

e) Polyamines

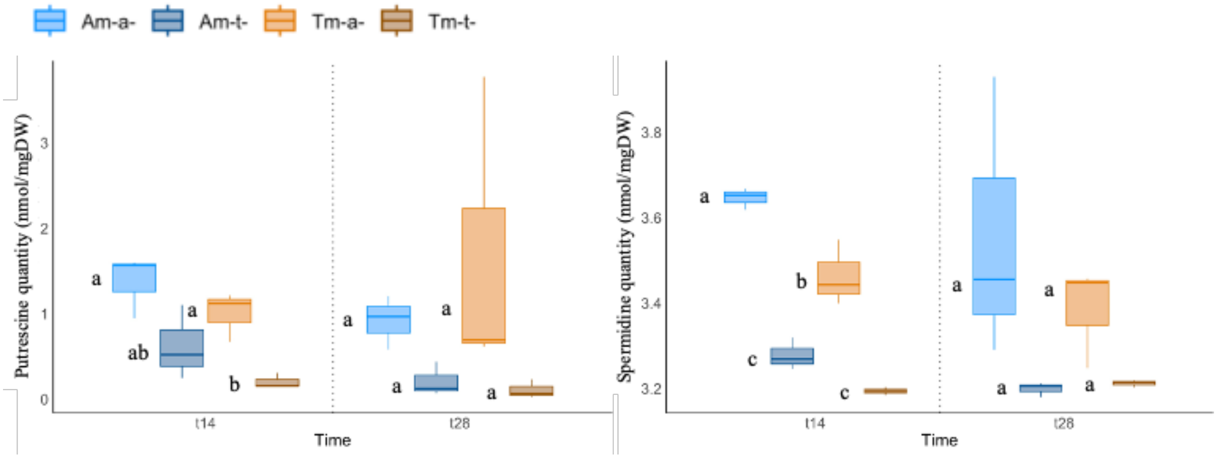


Others compounds

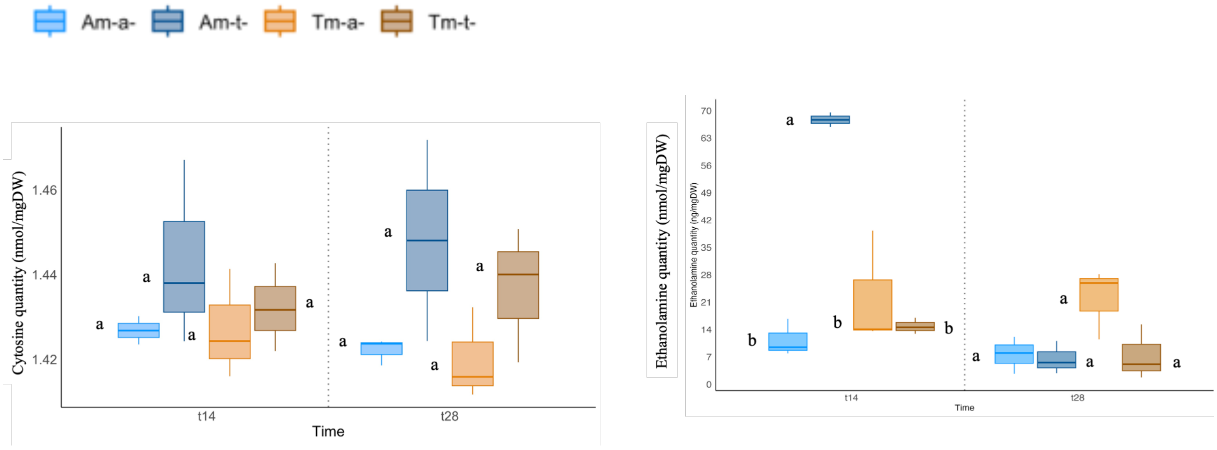


Figure S4: Continued; e) Polyamines and Others compounds.
