## Supplemental Table S1 for "Adaptive Strategies of the invasive aquatic plant, *Ludwigia grandiflora* subps. *hexapetala*: Contrasting Plasticity Between Aquatic and Terrestrial Morphotypes"

Table S1: ANOVA results for morphological, plant fitness, water content and allocation resources variables of aquatic and terrestrial morphotypes (Am and Tm) in both aquatic and terrestrial conditions (-a and -t) at 14 and 28 days. Significance codes: .001 "***"; .01 "**"; .05 "*"; nonsignificant NS"

| | **Analyses of traits (anova)** | | | | | | | | | | | | | | | | | | |
|---|---|---|---|---|---|---|---|---|---|---|---|---|---|---|---|---|---|---|---|
| | | | **Morphotype** | | | **Condition** | | | **Time** | | | **morphotype x condition** | | **morphotype x time** | | **condition x time** | | **Replicate** | |
| **Type of variable** | **Morphological trait** | **Data transformation** | Inequality | F value | P | Inequality | F value | P | Inequality | F value | P | F Value | P | F value | P | F value | P | F Value | P |
| **Morphological variables** | **LP** | BOXCOX | Am < Tm | 81.4428 | *** | a < t | 65.5728 | *** | t14 < t28 | 92.3161 | *** | 0.2814 | NS | 2.7244 | NS | 0.2133 | NS | 0.8299 | NS |
| | **nbL** | ORQ | Am < Tm | 60.3942 | *** | a<t | 133.5229 | *** | t14>t28 | 5.2899 | * | 2.2728 | NS | 13.7117 | *** | 78.2565 | *** | 18.3670 | *** |
| | **nbI** | BOXCOX | Am < Tm | 95.8488 | *** | a<t | 42.4311 | *** | t14 < t28 | 103.0203 | *** | 0.5102 | NS | 0.0005 | NS | 0.1894 | NS | 9.7614 | ** |
| | **nbNB** | Kruskall wallis (Chi_Square) | Am < Tm | 83.2930 | *** | a<t | 524.4279 | *** | t14=t28 | 3.2817 | NS | 46.1426 | *** | 11.5173 | *** | 2.4488 | NS | 2.7305 | NS |
| | **NbLLp** | BOXCOX | Am=Tm | 0.3982 | NS | a<t | 24.0978 | *** | t14>t28 | 154.6520 | *** | 6.7170 | * | 25.1848 | *** | 48.4830 | *** | 7.9201 | ** |
| | **NbILp** | BOXCOX | Am>Tm | 12.8168 | *** | a>t | 22.1517 | *** | t14>t28 | 16.0811 | *** | 0.8196 | NS | 4.0221 | * | 0.1294 | NS | 1.1654 | NS |
| **Plant fitness** | **DMT** | ORQ | Am<Tm | 77.3060 | *** | a<t | 527.2568 | *** | t14<t28 | 37.5683 | *** | 8.1808 | ** | 3.6327 | NS | 28.8053 | *** | 3.5180 | NS |
| **Water content** | **Sr** | ORQ | Am>Tm | 6.4058 | * | a>t | 378.0059 | *** | t14=t28 | 0.5036 | NS | 0.2350 | NS | 2.3859 | NS | 113.8392 | *** | 0.1980 | NS |
| | **Rr** | | Am=Tm | 0.0010 | NS | a < t | 57.8511 | *** | t14<t28 | 4.0632 | * | 0.4961 | NS | 0.9321 | NS | 8.3022 | ** | 0.3844 | NS |
| **Allocation ressources** | **Aloc S** | ORQ | Am=Tm | 2.6616 | NS | a=t | 3.8392 | NS | t14>t28 | 5.4438 | * | 3.9277 | * | 9.8596 | ** | 13.3122 | *** | 0.2502 | NS |
| | **Aloc R** | ORQ | Am=Tm | 2.6616 | NS | a=t | 3.8392 | NS | t14<t28 | 5.4438 | * | 3.9277 | * | 9.8596 | ** | 13.3122 | *** | 0.2502 | NS |
