## Supplemental Table S2-S6 for "Adaptive Strategies of the invasive aquatic plant, *Ludwigia grandiflora* subps. *hexapetala*: Contrasting Plasticity Between Aquatic and Terrestrial Morphotypes"

Table S2: Anova results for metabolite and phytohormone amounts of aquatic and terrestrial morphotypes (Am and Tm) in both aquatic and terrestrial conditions (a and t) at 14 days of experiment. Significance codes: .001 “***”; .01 “**”; .05 “*”; nonsignificant “NS”.

| | | **Analyses of metabolites quantities t14 (anova)** | | | | | | | |
|---|---|---|---|---|---|---|---|---|---|
| | **Factors** | **Morphotype** | | | **Condition** | | | **Morphotype x Condition** | |
| **Type of metabolites** | **Metabolites** | **inequality** | **F Value** | **P** | **inequality** | **F Value** | **P** | **F Value** | **P** |
| **Amino acids** | glycine | = | NS | NS | a>t | 7.232 | * | NS | NS |
| | isoleucine | = | NS | NS | a>t | 22.724 | ** | NS | NS |
| | leucine | = | NS | NS | = | NS | NS | 7.130 | * |
| | lysine | = | NS | NS | = | NS | NS | NS | NS |
| | methionine | = | NS | NS | = | NS | NS | NS | NS |
| | ornithine | = | NS | NS | = | NS | NS | NS | NS |
| | phenylalanine | = | NS | NS | a>t | 9.051 | * | NS | NS |
| | tyrosine | Am>Tm | 6.978 | * | = | NS | NS | NS | NS |
| | valine | = | NS | NS | a>t | 30.739 | ** | 13.315 | * |
| **Polyols** | adonitol | = | NS | NS | a>t | 56.755 | *** | 6.791 | * |
| | arabitol | = | NS | NS | = | NS | NS | 15.551 | ** |
| | erythritol | Am>Tm | 14.99 | ** | t>a | 17.70 | ** | 70.31 | *** |
| | glycerol | = | NS | NS | = | NS | NS | NS | NS |
| | G3P | = | NS | NS | a>t | 11.881 | * | NS | NS |
| | inositol | = | NS | NS | t>a | 6.050 | * | NS | NS |
| | mannitol | Am>Tm | 24.547 | ** | = | NS | NS | 57.987 | *** |
| **Sugars** | arabinose | Am>Tm | 162.163 | *** | t>a | 194.203 | *** | 162.489 | *** |
| | galactose | Am>Tm | 963.328 | *** | t>a | 959.586 | *** | 957.276 | *** |
| | glucose | Am>Tm | 63.253 | *** | t>a | 59.306 | *** | 54.598 | *** |
| | mannose | Am>Tm | 463.338 | *** | t>a | 550.840 | *** | 579.017 | *** |
| | ribose | Am>Tm | 170.155 | *** | t>a | 191.077 | *** | 169.010 | *** |
| | trehalose | Am>Tm | 58.292 | *** | = | NS | NS | NS | NS |
| | xylose | Am>Tm | 88.210 | *** | t>a | 133.169 | *** | 89.567 | *** |
| **Organic acids** | ascorbic acid | = | NS | NS | = | NS | NS | 7.152 | * |
| | gluconolactone | = | NS | NS | = | NS | NS | NS | NS |
| | glutamic acid | = | NS | NS | = | NS | NS | NS | NS |
| | glyceric acid | = | NS | NS | = | NS | NS | NS | NS |
| | malic acid | = | NS | NS | t>a | 22.496 | ** | NS | NS |
| | quinic acid | Tm>Am | 86.073 | *** | a>t | 190.063 | *** | 156.315 | *** |
| **Polyamines** | putrescine | Am>Tm | 6.347 | * | a>t | 24.487 | ** | NS | NS |
| | spermidine | Am>Tm | 28.618 | ** | a>t | 163.598 | *** | NS | NS |
| | spermine | = | NS | NS | = | NS | NS | NS | NS |
| **Others compounds** | cytosine | = | NS | NS | = | NS | NS | NS | NS |
| | ethanolamine | Am>Tm | 25.86 | ** | t>a | 35.15 | ** | 60.07 | *** |
| | Absisic acid | = | NS | NS | t>a | 107.393 | *** | NS | NS |

| | | | | | | | | | |
|---|---|---|---|---|---|---|---|---|---|
| **Phyto-hormones** | Auxin | = | NS | NS | = | NS | NS | 6.784 | * |
| | Salicylic acid | = | NS | NS | t>a | 9.006 | * | NS | NS |

Table S3: Anova results for metabolite and phytohormone amounts of aquatic and terrestrial morphotypes (Am and Tm) in both aquatic and terrestrial conditions (-a and -t) at 28 days of experiment. Significance codes: .001 “***”; .01 “**”; .05 “*”; nonsignificant “NS”. No significant effect was found for interaction between morphotype and condition.

| | | **Analyses of metabolites quantities t28 (anova)** | | | | | | | |
|---|---|---|---|---|---|---|---|---|---|
| | Factors | Morphotype | | | Condition | | | Morphotype x Condition | |
| **Type of metabolites** | Metabolites | inequality | F Value | P | inequality | F Value | P | F Value | P |
| | glycine | = | NS | NS | a>t | 46.327 | *** | NS | NS |
| | isoleucine | = | NS | NS | a>t | 29.159 | ** | NS | NS |
| | leucine | = | NS | NS | a>t | 15.745 | ** | NS | NS |
| | lysine | = | NS | NS | a>t | 6.950 | * | NS | NS |
| | methionine | = | NS | NS | a>t | 12.091 | * | NS | NS |
| | ornithine | = | NS | NS | a>t | 12.355 | * | NS | NS |
| | phenylalanine | = | NS | NS | a>t | 32.754 | ** | NS | NS |
| | tyrosine | = | NS | NS | a>t | 12.567 | * | NS | NS |
| | valine | = | NS | NS | a>t | 89.201 | *** | NS | NS |
| **Polyols** | adonitol | = | NS | NS | a>t | 14.746 | ** | NS | NS |
| | arabitol | = | NS | NS | a>t | 7.394 | * | NS | NS |
| | erythritol | = | NS | NS | = | NS | NS | NS | NS |
| | glycerol | = | NS | NS | a>t | 9.408 | * | NS | NS |
| | G3P | = | NS | NS | a>t | 20.483 | ** | NS | NS |
| | inositol | = | NS | NS | = | NS | NS | NS | NS |
| | mannitol | = | NS | NS | = | NS | NS | NS | NS |
| **Sugars** | arabinose | = | NS | NS | = | NS | NS | NS | NS |
| | galactose | Tm>Am | 6.386 | *** | = | NS | NS | NS | NS |
| | glucose | Tm>Am | 13.997 | ** | = | NS | NS | NS | NS |
| | mannose | Tm>Am | 6.054 | * | = | NS | NS | NS | NS |
| | ribose | = | NS | NS | = | NS | NS | NS | NS |
| | trehalose | = | NS | NS | = | NS | NS | NS | NS |
| | xylose | = | NS | NS | = | NS | NS | NS | NS |
| **Organic acids** | ascorbic acid | = | NS | NS | t>a | 17.171 | ** | NS | NS |
| | gluconolactone | = | NS | NS | t>a | 70.079 | *** | NS | NS |
| | glutamic acid | = | NS | NS | a>t | 12.336 | * | NS | NS |
| | glyceric acid | = | NS | NS | t>a | 6.847 | * | NS | NS |
| | malic acid | = | NS | NS | t>a | 8.112 | * | NS | NS |
| | quinic acid | = | NS | NS | = | NS | NS | NS | NS |
| **Polyamine** | putrescine | = | NS | NS | = | NS | NS | NS | NS |
| | spermidine | = | NS | NS | a>t | 8.341 | * | NS | NS |
| | spermine | = | NS | NS | a>t | 9.360 | * | NS | NS |

| | | | | | | | | | |
|---|---|---|---|---|---|---|---|---|---|
| **Others compounds** | cytosine | = | NS | NS | t>a | 9.728 | * | NS | NS |
| | ethanolamine | = | NS | NS | = | NS | NS | NS | NS |
| **Phytohormones** | Absissic acid | = | NS | NS | t>a | 107.393 | *** | NS | NS |
| | Auxin | = | NS | NS | = | NS | NS | 6.784 | * |
| | Salicylic acid | = | NS | NS | t>a | 9.006 | * | NS | NS |

**Table S4.** RDPI results and comparison of metabolites amounts of both morphotypes (Am and Tm) at 14-28 days. Significance codes: .001 "***"; .01 "**"; .05 "*"; non-significant (NS) "=" t-test. Phytohormone quantities weren't quantified at 14 days (shaded area). High plasticity index (bold), moderate plasticity index (bold italic), low plasticity index (normal). Inequality was indicated in blue when Am RDPI was superior to Tm RDPI and in orange when Tm RDPI was superior to Am RDPI.

| | | **RDPI analyses (t-test)** | | | | | | | |
|---|---|---|---|---|---|---|---|---|---|
| | **Temps** | **rdpi t14** | | | | **rdpi t28** | | | |
| Category | **Metabolites** | **Am** | **Tm** | **inequality** | **p** | **Am** | **Tm** | **inequality** | **p** |
| **Amino acid** | glycine | 0,24 | ***0,83*** | Tm>Am | *** | ***0,84*** | **0,91** | = | NS |
| | isoleucine | ***0,74*** | ***0,61*** | = | NS | ***0,88*** | ***0,77*** | Am>Tm | ** |
| | leucine | 0,34 | ***0,72*** | Tm>Am | *** | ***0,73*** | ***0,54*** | Am>Tm | * |
| | lysine | 0,34 | ***0,54*** | = | NS | 0,53 | ***0,83*** | Tm>Am | *** |
| | methionine | 0,14 | 0,16 | = | NS | 0,33 | 0,36 | = | NS |
| | ornithine | ***0,40*** | 0,08 | Am>Tm | ** | ***0,42*** | 0,25 | = | NS |
| | phenylalanine | ***0,62*** | ***0,50*** | = | NS | ***0,84*** | ***0,74*** | Am>Tm | * |
| | tyramine | 0,39 | 0,20 | = | NS | 0,09 | 0,22 | Tm>Am | * |
| | valine | 0,17 | ***0,72*** | Tm>Am | *** | ***0,76*** | ***0,68*** | = | NS |
| **Polyol** | Adonitol | 0,29 | ***0,62*** | Tm>Am | *** | ***0,48*** | ***0,36*** | = | NS |
| | erythritol | ***0,60*** | 0,32 | Am>Tm | *** | 0,28 | 0,22 | = | NS |
| | galactitol | ***0,45*** | 0,16 | Am>Tm | ** | 0,30 | 0,29 | = | NS |
| | G3P | 0,25 | 0,38 | Tm>Am | *** | ***0,52*** | ***0,63*** | = | NS |
| | inositol | ***0,54*** | 0,14 | Am>Tm | *** | 0,34 | 0,32 | = | NS |
| | sorbitol | 0,20 | 0,29 | = | NS | 0,11 | 0,19 | = | NS |
| | xylitol | 0,29 | 0,20 | = | NS | 0,13 | 0,18 | = | NS |
| **Sugar** | arabinose | **0,97** | 0,34 | Am>Tm | *** | ***0,53*** | ***0,62*** | = | NS |
| | galactose | **0,99** | 0,10 | Am>Tm | *** | ***0,67*** | 0,05 | Am>Tm | *** |
| | glucose | ***0,72*** | 0,06 | Am>Tm | *** | ***0,61*** | 0,20 | Am>Tm | *** |
| | mannose | **0,95** | 0,08 | Am>Tm | *** | ***0,45*** | 0,26 | = | NS |
| | ribose | **0,96** | 0,26 | Am>Tm | *** | 0,29 | 0,55 | = | NS |
| | trehalose | 0.15 | 0.10 | = | NS | 0,37 | 0,32 | = | NS |
| | xylose | **0,90** | ***0,40*** | Am>Tm | *** | ***0,65*** | ***0,47*** | = | NS |
| **Organic acid** | ascorbic acid | ***0,56*** | 0,26 | Am>Tm | ** | ***0,73*** | ***0,67*** | = | NS |
| | aspartic acid | ***0,57*** | 0,27 | Am>Tm | ** | 0,22 | 0,28 | = | NS |
| | citric acid | 0,30 | 0,13 | = | NS | 0,32 | ***0,40*** | = | NS |
| | gluconolactone | ***0,46*** | 0,31 | = | NS | ***0,53*** | ***0,67*** | = | NS |
| | lactic acid | 0,23 | ***0,59*** | Tm>Am | ** | 0,28 | ***0,63*** | Tm>Am | ** |
| | malic acid | ***0,51*** | 0,28 | Am>Tm | *** | ***0,42*** | 0,35 | = | NS |
| | quinic acid | 0,17 | ***0,76*** | Tm>Am | *** | 0,28 | 0,32 | = | NS |
| | succinic acid | ***0,46*** | 0,18 | Am>Tm | ** | ***0,40*** | ***0,46*** | = | NS |
| **Polyamine** | putrescine | ***0,42*** | ***0,65*** | Tm>Am | * | ***0,64*** | ***0,81*** | = | NS |
| | spermidine | 0,05 | 0,04 | Am>Tm | ** | 0,05 | 0,03 | = | NS |
| **Other** | dopamine | ***0,53*** | 0,18 | Am>Tm | ** | 0,30 | 0,26 | = | NS |
| | ethanolamine | ***0,72*** | 0,19 | Am>Tm | *** | 0,31 | ***0,55*** | = | NS |
| | octopamine | 0,06 | 0,03 | = | NS | ***0,41*** | 0,06 | Am>Tm | ** |
| **Phytohormone** | abssicic acid | | | | | ***0.75*** | ***0.70*** | = | NS |
| | Auxin | | | | | ***0.50*** | 0.20 | Am>Tm | *** |
| | Salicylic acid | | | | | ***0.47*** | ***0.50*** | = | NS |

**Table S5.** RDPI results and comparison of metabolites amounts of both morphotypes (Am and Tm) showing the dynamic of rdpi during time. Significance codes: .001 "***"; .01 "**"; .05 "*"; non-significant (NS) "=" t-test. Inequality of RDPI dynamic was indicated in yellow when t14 value was superior to t28 value and in green when t28 was superior to t14.

| | | RDPI analyses (t-test) | | | | | | | |
|---|---|---|---|---|---|---|---|---|---|
| | Temps | rdpi dynamics Am | | | | rdpi dynamics tm | | | |
| Category | Metabolites | t14 | t28 | inequality | p | t14 | t28 | inequality | p |
| **Amino acid** | glycine | 0.24 | 0.84 | t28>t14 | *** | 0,83 | 0,91 | t28>t14 | ** |
| | isoleucine | 0.73 | 0.88 | = | NS | 0,61 | 0,77 | = | NS |
| | leucine | 0.34 | 0.73 | t28>t14 | *** | 0,72 | 0,54 | t14>t28 | * |
| | lysine | 0.34 | 0.53 | = | NS | 0,54 | 0,83 | t28>t14 | * |
| | methionine | 0.14 | 0.32 | t28>t14 | * | 0,16 | 0,36 | t28>t14 | ** |
| | ornithine | 0.20 | 0.15 | = | NS | 0,08 | 0,25 | t28>t14 | * |
| | phenylalanine | 0.62 | 0.84 | t28>t14 | ** | 0,50 | 0,74 | t28>t14 | * |
| | tyramine | 0.39 | 0.09 | t14>t28 | *** | 0,20 | 0,22 | = | NS |
| | valine | 0.17 | 0.76 | t28>t14 | *** | 0,72 | 0,68 | = | NS |
| **Polyol** | Adonitol | 0.29 | 0.48 | t28>t14 | * | 0,62 | 0,36 | t14>t28 | ** |
| | erythritol | 0.60 | 0.28 | t14>t28 | *** | 0,32 | 0,22 | = | NS |
| | galactitol | 0.45 | 0.30 | = | NS | 0,16 | 0,29 | = | NS |
| | G3P | 0.24 | 0.52 | t28>t14 | ** | 0,38 | 0,63 | t28>t14 | *** |
| | inositol | 0.55 | 0.34 | = | NS | 0,14 | 0,32 | t28>t14 | * |
| | sorbitol | 0.20 | 0.11 | = | NS | 0,29 | 0,19 | t14>t28 | * |
| | xylitol | 0.29 | 0.13 | t14>t28 | * | 0,20 | 0,18 | = | NS |
| **Sugar** | arabinose | 0.96 | 0.53 | t14>t28 | *** | 0,34 | 0,62 | = | NS |
| | galactose | 0.99 | 0.67 | t14>t28 | *** | 0,10 | 0,05 | = | NS |
| | glucose | 0,72 | 0,61 | = | NS | 0,06 | 0,20 | t28>t14 | * |
| | mannose | 0,95 | 0,45 | t14>t28 | *** | 0,08 | 0,26 | t28>t14 | * |
| | ribose | 0,96 | 0,29 | t14>t28 | *** | 0,26 | 0,55 | = | NS |
| | trehalose | 0.15 | 0.37 | = | NS | 0.10 | 0,32 | t28>t14 | * |
| | xylose | 0,90 | 0,65 | t14>t28 | ** | 0,40 | 0,47 | = | NS |
| **Organic acid** | ascorbic acid | 0,56 | 0,73 | t28>t14 | * | 0,26 | 0,67 | t28>t14 | *** |
| | aspartic acid | 0,57 | 0,22 | t14>t28 | *** | 0,27 | 0,28 | = | NS |
| | citric acid | 0,30 | 0,32 | = | NS | 0,13 | 0,40 | t28>t14 | *** |
| | gluconolactone | 0,46 | 0,53 | = | NS | 0,31 | 0,67 | t28>t14 | *** |
| | lactic acid | 0,23 | 0,28 | = | NS | 0,59 | 0,63 | = | NS |
| | malic acid | 0,51 | 0,42 | = | NS | 0,28 | 0,35 | = | NS |
| | quinic acid | 0,17 | 0,28 | = | NS | 0,76 | 0,32 | t14>t28 | *** |
| | succinic acid | 0,46 | 0,40 | = | NS | 0,18 | 0,46 | t28>t14 | ** |
| **Polyamine** | putrescine | 0,42 | 0,65 | = | NS | 0,65 | 0,81 | = | NS |
| | spermidine | 0,05 | 0,04 | = | NS | 0,04 | 0,03 | t14>t28 | * |
| **Other** | dopamine | 0,53 | 0,30 | t14>t28 | * | 0,18 | 0,26 | = | NS |
| | ethanolamine | 0,72 | 0,19 | t14>t28 | *** | 0,19 | 0,55 | t28>t14 | ** |
| | octopamine | 0,06 | 0,41 | t28>t14 | ** | 0,03 | 0,06 | = | NS |

**Table S6**: Metabolomic pathways identified by metaboanalyst analysis in shoots in both aquatic and terrestrial conditions (a and t) at 14 (t14) and/or 28 (t28) days of experiment. Results are based on comparison between conditions (a *vs* t), p-holm<0.05.

| Time | Condition | Metabolomic pathways | List of metabolites involved |
|---|---|---|---|
| t14 and t28 | a>t | Valine, leucine, isoleucine degradation pathway | Isoleucine, leucine and valine |
| | | Valine, leucine, isoleucine biosynthesis pathway | Isoleucine, leucine, valine |
| | | Glucosinolate biosynthesis pathway | Isoleucine, leucine, methionine, phenylalanine and valine |
| | | Aminoacyl-tRNA biosynthesis pathway | Alanine, asparagine, aspartic acid, glutamic acid, glycine, isoleucine, lysine, methionine, phenylalanine, serine, threonine, tyrosine and valine |
| t28 | a>t | Phenylalanine metabolism pathway | Phenylalanine |
| | | Phenylpropanoide biosynthesis pathway | Phenylalanine |
| | | Phenylalanine, tyrosine and tryptophan biosynthesis pathway | Phenylalanine and tyrosine |
